# Can we use it? On the utility of *de novo* and reference-based assembly of Nanopore data for plant plastome sequencing

**DOI:** 10.1101/855981

**Authors:** Agnes Scheunert, Marco Dorfner, Thomas Lingl, Christoph Oberprieler

## Abstract

The chloroplast genome harbors plenty of valuable information for phylogenetic research. Illumina short-read data is generally used for *de novo* assembly of whole plastomes. PacBio or Oxford Nanopore long reads are additionally employed in hybrid approaches to enable assembly across the highly similar inverted repeats of a chloroplast genome. Unlike for PacBio, plastome assemblies based solely on Nanopore reads are rarely found, due to their high error rate and non-random error profile. However, the actual quality decline connected to their use has never been quantified. Furthermore, no study has employed reference-based assembly using Nanopore reads, which is common with Illumina data. Using *Leucanthemum* Mill. as an example, we compared the sequence quality of seven plastome assemblies of the same species, using combinations of two sequencing platforms and three analysis pipelines. In addition, we assessed the factors which might influence Nanopore assembly quality during sequence generation and bioinformatic processing.

The consensus sequence derived from *de novo* assembly of Nanopore data had a sequence identity of 99.59% compared to Illumina short-read *de novo* assembly. Most of the found errors comprise indels (81.5%), and a large majority of them is part of homopolymer regions. The quality of reference-based assembly is heavily dependent upon the choice of a close-enough reference. Using a reference with 0.83% sequence divergence from the studied species, mapping of Nanopore reads results in a consensus comparable to that from Nanopore *de novo* assembly, and of only slightly inferior quality compared to a reference-based assembly with Illumina data (0.49% and 0.26% divergence from Illumina *de novo*). For optimal assembly of Nanopore data, appropriate filtering of contaminants and chimeric sequences, as well as employing moderate read coverage, is essential.

Based on these results, we conclude that Nanopore long reads are a suitable alternative to Illumina short reads in plastome phylogenomics. Only few errors remain in the finalized assembly, which can be easily masked in phylogenetic analyses without loss in analytical accuracy. The easily applicable and cost-effective technology might warrant more attention by researchers dealing with plant chloroplast genomes.

## 1. Introduction

The chloroplast (cp) is the most characteristic organelle of plant cells. It carries its own unique genome, whose genes are mainly related to housekeeping functions and photosynthesis [1, 2]. Most cp genomes are highly conserved regarding structure, size, and functionality of their genes. Their molecular conformation has long been thought to be exclusively circular, but growing evidence suggests that plastid chromosomes often form concatemers of several molecules and can occur in circularized and/or linear form [3]; for further references see [1]. One genome equivalent has a quadripartite structure consisting of two regions of unique DNA termed the large and small single-copy region (LSC and SSC, respectively), and a pair of near-identical large inverted repeats (IR_B_ and IR_A_) situated in between them [4]. Low substitution rates [5], very low levels of recombination, and high copy numbers in cells make the haploid genomes an easily accessible source of valuable phylogenetic information [6], for example by Sanger sequencing of molecular markers following PCR. Since the advent of next-generation sequencing (NGS) techniques, a growing number of research projects has taken advantage of the possibility to build phylogenetic analyses upon sequences of whole cp genomes instead of only a few molecular markers. In consequence, as of 19/10/2019, 2,982 angiosperm chloroplast genomes have been available in NCBI’s Genbank (https://www.ncbi.nlm.nih.gov/genbank/). Apart from a substantially higher number of informative characters, larger-scale features like major inversions and gene losses can now be studied in detail.

The standard method applied for plastome recovery is Illumina short-read sequencing and assembly. The reads are of high quality (app. 0.3% error rate for the Illumina MiSeq [7]), but currently limited to 300 basepairs (bp) paired-end reads. This means that a cp genome containing two IRs cannot be reliably assembled into one continuous contig, as repeat regions can only be resolved up to the length of the sequenced insert size. By contrast, third-generation single-molecule sequencing yields reads of much higher lengths: with PacBio SMRT DNA sequencing, average read length is now up to 20 kb, (according to Pacific Biosciences website at https://www.pacb.com/products-and-services/sequel-system/latest-system-release/, accessed 30/10/2019). The alternative provided by Oxford Nanopore MinION features average read lengths of more than 6-8 kb, with also more than 20 kb regularly achieved depending on input DNA quality; and maximum read lengths exceeding 150 kb [8]. The main problem with these techniques is their high initial error rate. PacBio raw reads have an error rate of 11-15%; circular consensus sequencing can lower this value to < 1% [9], however at the cost of reduced sequencing depth and/or multiplexing capacity. Nanopore reads, despite several improvements, still feature a raw error rate of 5-15% [10]. This problem can be alleviated by the use of consensus techniques [via Partial Order Alignment (POA) graphs]. Loman et al. (2015) integrated read correction via POA graphs into an assembly pipeline, thus increasing nucleotide identity to a reference genome from initially 80.5% to 97.7% in the reads [11]. The authors furthermore introduced post-assembly processing with Nanopolish, and finally obtained an assembly with 99.5% nucleotide identity. However, in contrast to the random error profile in PacBio, some errors in Nanopore reads seem to be biased: A/T substitution errors are less likely than all other substitution errors, and deletion errors, which are the most common errors found in Nanopore reads, increase in homopolymer runs [7] (see also [12]). This means that a remaining 0.5% error rate (see above) cannot be filtered using consensus techniques because they are essentially non-random.

Long reads from both sequencing platforms have been introduced into so-called hybrid approaches, which intend to combine the advantages of long, possibly repeat-spanning reads (Nanopore, PacBio) with the low error rate of short (paired-end) Illumina sequences to assemble nuclear genomes [13, 14] and complete cp genomes [15, 16]. Regarding non-hybrid assembly, PacBio has been used for the sequencing of whole microbial genomes (e.g., [17]; with many following since). Full plastomes have also been reported, among others by [18, 19] or Ferrarini et al. (2013) [20], who also disambiguated the two IR copies. However, assembly yielding a single full contig without any necessary post-assembly processing still seems to be problematic as pointed out by [21].

Exclusively using Nanopore data for genome assembly is less widespread, but nevertheless has led to a growing number of sequenced organisms, especially microbes (see [22]), but also larger genomes (e.g., [23]). However, to our knowledge, only one study has relied solely upon Nanopore data for plastome assembly up to now [24]. Additionally, Wang et al. (2018) presented a comparison of the techniques for assembling cp genomes, via short reads, long Nanopore reads or a hybrid approach involving both [21]. Furthermore, the effect of varying read coverages of long and short-read data was also examined. But no study has yet aimed to quantify the loss in assembly quality that is linked to the exclusive use of Nanopore reads during plastome sequencing. Furthermore, assembly of Nanopore sequence data is routinely done by *de novo* approaches. No one has as yet assessed the quality of plastome sequences derived from reference-based assembly, although mapping assembly of plastomes is common with Illumina data. If plastome sequences assembled from Nanopore data showed only minor declines compared to the present “gold standard” Illumina *de novo* assembly, Nanopore could be a reasonable alternative to Illumina or PacBio sequencing for several research questions. Also, this would mean lower sequencing costs, as only a comparatively small amount of sequence data is needed for assembly of a complete plastome and many individuals thus can be multiplexed on a single MinION flow cell. Additionally, unlike Illumina or PacBio, the device is low-priced and easily installed in an average lab.

The aims of the present study were therefore: 1) to compare the sequence quality of a plastome derived from Nanopore data *de novo* assembly to a corresponding assembly based on Illumina data, with the goal to quantitate their differences on the basepair level; 2) to explore the potential of reference-based assembly using Nanopore data and to evaluate the quality of these assemblies with respect to Nanopore or Illumina *de novo* assemblies; 3) to categorize the error types associated with Nanopore-derived plastome sequences and to assess if and how these errors and possible biases in their distribution might impact the use of Nanopore-only cp genomes for phylogenomic research; and 4) to provide practical guidelines for both *de novo* and mapping assembly of Nanopore data for researchers interested in establishing Nanopore sequencing in their lab.

To this end, we here present complete cp genomes from two species of the genus *Leucanthemum* Mill., which belongs to subtribe Leucantheminae in the Compositae tribe Anthemideae. The provided plastomes are the first for the subtribe. The chosen study group forms a polyploid complex of c. 42 often closely related species with a main distribution in Europe and the Mediterranean. The plastid genomes of the two species [*L. vulgare* Lam. and *L. virgatum* (Desr.) Clos] were generated by long-range PCR combined with *de novo* assembly of Illumina MiSeq short reads. Both were examined regarding their sequence divergence and surveyed for suitable new markers for infrageneric phylogenetic analyses in *Leucanthemum*. Subsequently, the *L. vulgare* plastome served as “gold standard” reference for comparisons of different assembly methods and types of data. To achieve this, the cp genome sequence was re-generated six times: first, reference-based assembly of the short reads was performed, with a closely (*L. virgatum*) as well as a more distantly related taxon (*Artemisia frigida* Willd., belonging to subtribe Artemisiinae of tribe Anthemideae [25]) as reference for mapping, to assess the influence of reference choice on assembly quality. *De novo* and reference-based assembly approaches were then repeated using long-read Nanopore data instead of Illumina short reads. In a third step, we tested whether a correction of the error-prone Nanopore raw reads by high-quality Illumina reads improves the results of a subsequent Nanopore *de novo* assembly (hybrid approach).

## 2. Materials and Methods

We provide a detailed account on the methods, tools and parameters employed in the present study, as correct setting of suitable options is essential for obtaining good results in several cases. A schematic workflow is available in Fig 1.

**Fig 1.**
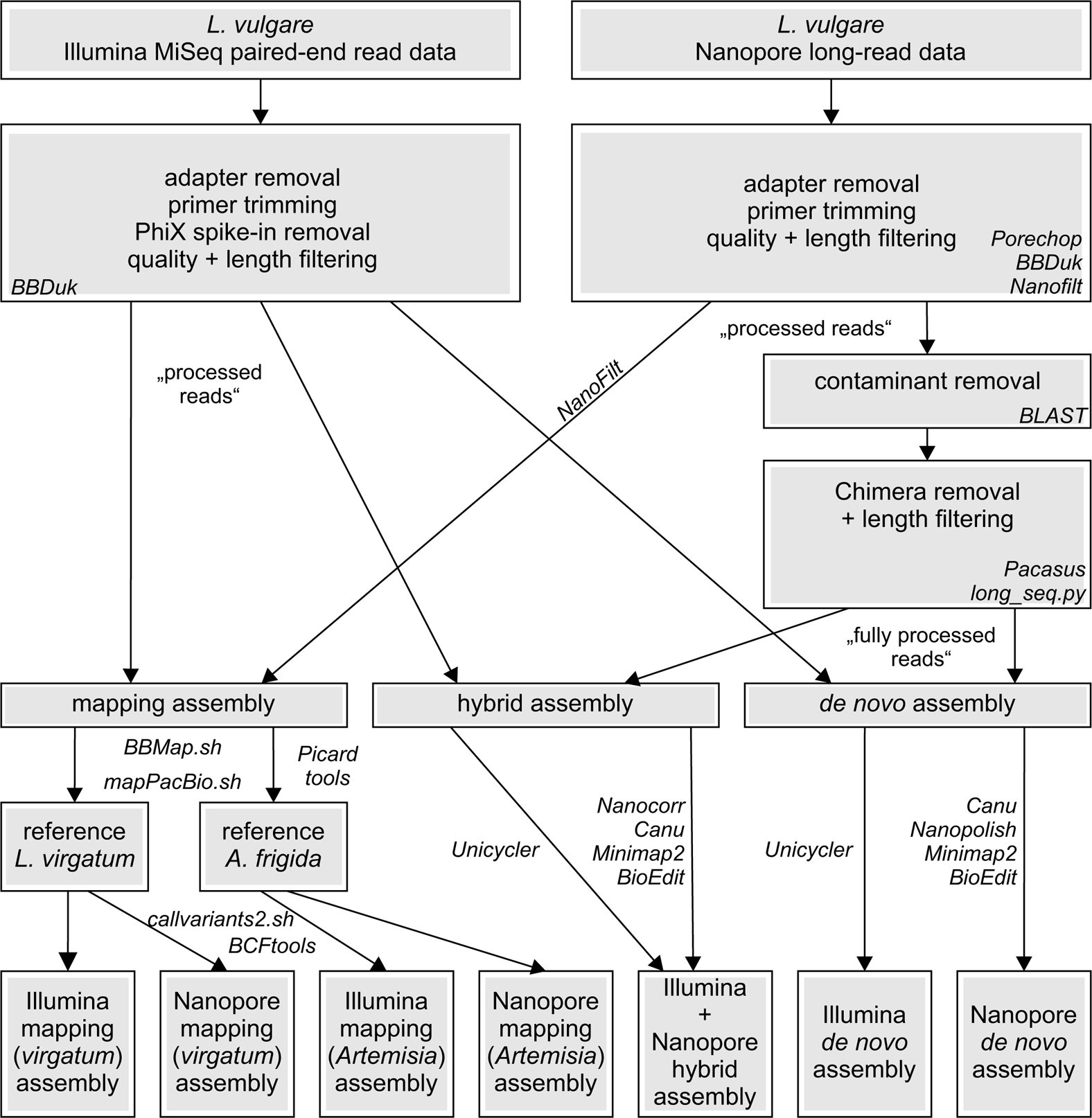
Schematic bioinformatic workflow for comparative assembly of chloroplast genomes. Tools used for each processing step are given in italics. *Leucanthemum virgatum* was assembled *de novo* with Illumina data and subsequently used as reference for mapping.

### 2.1. Plant material and DNA extraction

Silica-gel dried leaf tissue from two accessions of *L. vulgare* and *L. virgatum*, collected during field excursions, was used for the present study (for voucher details see Supporting Information S1 Table). Genomic DNA (gDNA) was extracted following a CTAB extraction protocol adapted from [26] and [27]. Each extraction was quality-checked via agarose gel electrophoresis, and DNA concentration and purity was determined using a NanoDrop spectrophotometer One (Thermo Fisher Scientific, Waltham MA, USA).

### 2.2. Amplification via long-range PCR

Chloroplast DNA was amplified using 32 primer pairs by [28] to generate 16 overlapping PCR fragments spanning the whole chloroplast. Before amplification, we assessed their potential in Anthemideae by fitting them onto *Artemisia frigida* (available with the ID NC_020607.1 in Genbank). This was done by performing megaBLAST searches [29] at the National Center for Biotechnology Information (NCBI) BLAST website https://blast.ncbi.nlm.nih.gov/Blast.cgi [30]. Based on the estimated product sizes and overlap lengths in *A. frigida*, for fragments 2-5, other primers were chosen (and sometimes slightly modified) from the list of primers given in [28]; these fragments span the region of the LSC harboring the two inversions which are characteristic for Asteraceae [31, 32], but are not found in Orobanchaceae [28]. Furthermore, primer ndhI.194R (fragment 15) was replaced by the reverse-complement of primer ndhA.535F (fragment 14) due to amplification problems, and several primers were slightly modified. All used primer sequences are available in Supporting Information S2 Table. After these modifications, estimated fragment sizes in *Artemisia* varied between ∼6,000 and 12,300 bp, with a minimum overlap of 173 bp among fragments, except for a small sequence gap between fragment 14 and 15 (due to the double use of primer ndhA.535F), which was filled by additional Sanger sequencing (see below). PCR reactions were performed using the Invitrogen Platinum SuperFi Green PCR Master Mix (Thermo Fisher Scientific). Amplicons were purified either using an ethanol precipitation protocol modified from [33] with further optimizations, or with the AmpliClean^TM^ Magnetic Bead-based PCR cleanup (Nimagen, Nijmegen, Netherlands), and were checked by gel electrophoresis before and after purification. DNA concentration and purity was again determined via Nanodrop and a Qubit 2.0 fluorometer (Thermo Fisher Scientific). All 16 fragments were then equimolarily pooled for each sample and subjected to library preparation procedures.

### 2.3. Illumina sequencing

Four libraries were generated for Illumina sequencing, two each for *L. vulgare* and *L. virgatum*, with different fragment sizes for two MiSeq runs of 300 bp and 80 bp paired-end reads, respectively (see below). Library preparation was done using the NEBNext Ultra II FS DNA Library Prep Kit for Illumina (New England BioLabs, Frankfurt, Germany) and the protocol for use with inputs of ≥ 100 ng DNA. 5 min (*L. vulgare*) and 10 min (*L. virgatum*) of enzymatic fragmentation produced fragments of around 400 bp; 20 min (*L. vulgare*) and 35 min (*L. virgatum*) of fragmentation were necessary to obtain ∼200-bp fragments. As post-ligation size selection following the manufacturer’s protocol did not work, a two-step selection using AMPure XP beads was developed, where, for the 300 bp run, 0.7x beads to DNA volume will remove fragments below 400 bp in a first step, and re-suspending the eluate DNA with 0.55x beads will keep fragments of > 600 bp bound to the beads in a second step, the supernatant hence containing fragments of the desired length (here around 500 bp). For the 80 bp run, a larger fragment range was obtained using a similar approach, by first binding and discarding fragments larger than 500 bp using 0.65x beads and then concentrating the supernatant with 1.2x beads, which will bind everything except unwanted very small molecular weight DNA. The eluate thus consisted of fragments of c. 100-500 bp length, with a peak of c. 330 bp. As the PCR enrichment of adapter-ligated DNA preferentially enriched the smaller fragments, the final libraries had a peak length of around 300 bp for the 300 bp run and c. 220 bp for the 80 bp run. The 6-bp NEBNext indices 6 and 12 were used for barcoding *L. vulgare* and *L. virgatum*, respectively. Samples were evaluated on an Agilent 4200 TapeStation using the High Sensitivity D1000 ScreenTape assay (Agilent Technologies, Waldbronn, Germany). For the 300-bp run, the two libraries then were pooled with 48 libraries of other projects and subsequently sequenced on 0.09% (*L. virgatum*) and 0.08% (*L. vulgare*) of one lane on an Illumina MiSeq system (Illumina, San Diego, California, USA) at the Department of Biochemistry I at the University of Regensburg. The 80-bp run was conducted on the same sequencer; *L. virgatum* was sequenced on 1.13% and *L. vulgare* on 1.11% of one lane, together with 22 other libraries.

### 2.4. Nanopore sequencing

A Nanopore sequencing library was generated for *L. vulgare* using the 1D Native barcoding genomic DNA protocol, with the PCR-free SQK-LSK108 Ligation Sequencing kit and EXP-NBD103 Native Barcoding Expansion 1-12. Library preparation was done according to the manufacturer’s protocol except for some minor modifications; also, no DNA fragmentation and no FFPE DNA repair was performed. As the library contained one other sample, *L. vulgare* was barcoded with the 24-bp barcode NB 07. Sequencing was performed using a FLO-MIN106 flow cell on a Nanopore MinION sequencing device (Oxford Nanopore Technologies, Oxford, UK) at the Department of Microbiology at the University of Regensburg.

### 2.5. Complementary Sanger sequencing

For exact determination of the LSC/IR/SSC boundaries, primers were designed spanning the four junctions (Supporting Information S2 Table) and amplified PCR fragments were then Sanger sequenced. Likewise, primers were designed to span the gap between long-range PCR fragments 14 and 15. Sanger sequences from this gap region were used to correct and/or complete all eight assembled consensus sequences generated in the present study.

### 2.6. Processing of Illumina reads and genome assembly

#### 2.6.1. Read trimming and filtering

Basic read statistics for Illumina and Nanopore data were obtained using the GenomeTools v.1.5.10 [34] tool *seqstat*. Regarding Illumina data, base-call files from both MiSeq runs were converted into fastq format read files, and reads were demultiplexed using bcl2fastq v.2.19 (Illumina). For each sample, reads from both runs were merged into one dataset. The paired-end read mates of each *L. vulgare* and *L. virgatum* were then processed using BBDuk from the BBTools v.38.29 software suite [35]: adapter and barcode sequences as well as all downstream basepairs were clipped from the reads, PhiX contaminant reads were removed, and primer sequences deriving from the long-range PCRs were trimmed sequentially from 5’- and 3’-ends of reads. This is important as primer sequences need not necessarily match the true sequence of the sample and thus might introduce wrong signal into an assembly. Afterwards, paired-end reads were subjected to quality trimming and length filtering also via BBDuk, keeping only reads composed of regions with Phred quality scores of Q = 20 or higher, and of minimum length = 50 bp. Trimming and filtering success was verified by examining the fastq files using FastQC v.0.11.8 [36]. Processed reads were used as input for *de novo* as well as mapping assemblies.

#### 2.6.2. Reference-based mapping assembly of Illumina data

Reference-based assembly of *L. vulgare* reads used two different references (see also section 2.7.2), the sequence from the Illumina *de novo* assembly of *L. virgatum* and the chloroplast genome of *Artemisia frigida* [further referred to as “mapping (*virgatum*)” and “mapping (*Artemisia*)”]. The gapped read aligner BBMap (*bbmap.sh* in BBTools software suite v.38.58), which also provides several mapping statistics, was chosen for mapping paired-end reads in order to accommodate larger indels possibly present between the two genomes. BBMap was run in very slow mode (*vslow = t*); the maximum indel size to search for was set to 300 bp for mapping to *L. virgatum* and to 1,500 bp for *A. frigida*. Unpaired reads were considered unmapped (*pairedonly = t*), *killbadpairs* set to *true* and *pairlen* to 180 bp. If a read had more than one top-scoring mapping location, one site was selected randomly (*ambiguous = random*), so that the resulting .sam files did not contain secondary mappings.

The .sam files were converted to .bam format, sorted and indexed with SAMtools v.1.9 [37]. For removal of PCR and optical duplicate reads in the dataset, Picard tools *MarkDuplicates* v.2.16.0 (Broad Institute of MIT and Harvard, Cambridge, Massachusetts, USA; available at http://broadinstitute.github.io/picard) was used. Mappings were evaluated regarding contiguity, coverage, and mapping quality with Qualimap v.2.2.1 [38] and the Integrative Genomics Viewer (IGV) v.2.4.15 [39]. For generating a consensus from the mapped read data, variants were then filtered and called with the shell script *callvariants2.sh* from the BBTools software suite (v.38.32). The script was set to ignore alignments with mapping quality lower than one (*minreadmapq = 1*). Setting a low value here is important as otherwise, variants from those parts of the IR which are represented twice in the reference, would not be called. Furthermore, no realignment and no trimming were performed and *ploidy* was set to 1. *Rarity* and *minallelefraction* parameters were set to 0.51, which penalizes and ignores variants with lower allele frequencies, enabling a “majority vote” among present variants as well as the reference base. All other variant filters were kept on default values. The resulting .vcf files were subsequently compressed and indexed with *bgzip* and *tabix* from the HTSlib package v.1.9 [37]. Variant statistics were calculated using VCFtools v.0.1.15 [40]. The consensus sequences were generated from the variants using the BCFtools [37] command *bcftools consensus*, setting missing genotypes to be represented by Ns instead of skipping them.

#### 2.6.3. *De novo* assembly of Illumina data

*De novo* assembly of *L. vulgare* and *L. virgatum* reads was done with the Unicycler pipeline v.0.4.7 [41], which mainly relies on the de Bruijn graph-based *de novo* assembler SPAdes v.3.13.0 [42] and the assembly polishing tool Pilon v.1.22 [43]. Running the pipeline in *conservative* mode ensured the lowest possible misassembly rate. As a result, contigs linked in a circularized assembly graph were obtained, which was visualized and assessed with Bandage v.0.8.1 [44]. Ambiguities observed in the graph are due to the presence of the large IR in the plastome. Fragments with multiplicity > 1.0 were assumed to belong to the IR; this was additionally confirmed by megaBLAST website searches. Ambiguities were then resolved in Bandage, by doubling the contigs associated with the IR to establish unique connections between contigs. The finalized chloroplast genome sequence could be extracted in fasta format. As *L. virgatum* was assembled with the SSC in the alternate orientation with respect to *L. vulgare*, the SSC was reverse-complemented for all subsequent mapping and comparative analyses. For obtaining read coverages, the input reads used in the *de novo* assemblies were mapped to the assembled cp genome sequences without the IR_A_. Mapping was done with BBMap as outlined in section 2.6.2., however with *maxindel* lowered to 100.

### 2.7. Processing of Nanopore reads and genome assembly

#### 2.7.1. Base calling, read trimming, and filtering

Base calling was done using Albacore v.2.3.4 (Oxford Nanopore Technologies). Base-called reads were quality-sorted by the software, with only reads with a mean Q-score > 7 put to a “pass” folder, and demultiplexed according to all found barcodes. Sequencing statistics were summarized using ShinyNANO v.0.1 (at www.shinynano.com). Nanopore adapter and barcode sequences were trimmed from the reads by submitting the entire pass folder to Porechop v.0.2.4 (available at https://github.com/rrwick/Porechop), with the *--discard_middle* option turned on and *--middle_threshold* set to 75. This will discard reads containing a ≥ 75% adapter-match in their middle, which might indicate a chimera. Porechop also re-demultiplexed the reads, and only those with barcode NB 07 recognized by both Porechop and Albacore were used for further analyses. Primer trimming was done sequentially at the 5’- and 3’-end of reads, using BBDuk (BBTools software suite v.38.29) similar to primer trimming for Illumina data, with the following settings to accomodate the high error rate of Nanopore sequence data, while assuring that only real primer sequence was trimmed: *ktrim= l/r*, *k = 11*, *hdist = 0*, *edist = 2*, *mm = f*, *rcomp = f*, *mkf = 0.51*, *restrictleft/restrictright = 100*, and *copyundefined = t*. As references, primer sequences were given in normal, complement, reverse, and reverse-complement orientations. Trimmed reads were then subjected to a length- and quality-filtering step in NanoFilt v.2.2.0 [45], leaving only reads with lengths ≤ 12,600 bp and a minimum average read Q-score of 7 in the datasets. Raw reads as well as trimmed and quality- and length-filtered reads (“processed reads”) were examined using FastQC. As base quality values were to be preserved for reference-based mapping assembly, no further processing of reads was done for this approach.

For *de novo* assembly, reads were further optimized: possible contaminant reads were filtered by doing a local megaBLAST search against the *Artemisia frigida* chloroplast genome (with the IR_A_ removed) with BLAST+ v.2.7.1 [46] and using the BLAST output for a filtering step, in which only reads with one or more hits on the *Artemisia* genome over at least 70% of their length were kept. Furthermore, to split possible chimeric reads resulting from erroneous amplification during long-range PCR, Pacasus v.1.1.1 [47] was used with the following settings: *--minimum_read_length = 0*, *--filter_factor = 0.0001*, *--query_coverage = 0.001*, *--query_identity = 0.001*, *--relative_score = 0.001*, and *--base_score = 0.50*. As no length filter was applied, very short sequences with lengths < 50 bp were removed afterwards in an additional step using the python script *long_seq.py* (available at http://seqanswers.com/forums/showthread.php?t=31046). The finished, trimmed, quality- and length-filtered, decontaminated, and putatively chimera-free reads are referred to as “fully processed reads” hereafter.

#### 2.7.2. Reference-based mapping assembly of Nanopore data

Processed reads from *L. vulgare* were analyzed to evaluate the suitability of the mapping approach for Nanopore data. To avoid problems caused by too high read coverage (see Discussion, section 4.6.), a reduced dataset similar to that employed for *de novo* assembly (see below) was used: only reads with a minimum length of 4,200 bp were kept after another filtering step with NanoFilt executed with option *-l 4200*. References for mapping were identical to those from the mapping assembly of Illumina data. Importantly, to avoid secondary mappings, but also mismappings and subsequent false variant calls as far as possible, the references were designed to contain only those parts of the second inverted repeat (IR_A_) needed to assure correct mapping of long reads from fragments 15 and 16, which span the borders between the SSC and the IR_A_, and the IR_A_ and the LSC, respectively. Mapping was done using the *mapPacBio.sh* script (BBTools software suite v. 38.58), with settings analogous to those employed for Illumina data in *bbmap.sh* regarding the *vslow*, *ambiguous*, and *maxindel* parameters. Additionally, as the script only correctly processes reads up to a certain length, the long Nanopore reads were split into 3,000-bp chunks using the *maxlen = 3000* option. Processing and evaluation of the resulting mappings also followed procedures described in section 2.6.2., except for the removal of duplicates, as Nanopore reads are derived from unfragmented PCR products, and hence were expected to feature high portions of identical starting positions in the mapping. Variant calling again was done with *callvariants2.sh*. Settings for *ploidy* and *minreadmapq* corresponded to those for Illumina data, and no trimming was performed. However, to improve mapping accuracy around indels (which are expected to occur more often when mapping Nanopore data), realignment was performed using settings *realign = t*, *repadding = 70*, *rerows = 3000* (chosen according to input read length), *recols = 3740*, and *msa = MultiStateAligner9PacBio*. Variant filtering parameters were chosen as follows: *minreads = 2*, *minqualitymax = 10*, *minedistmax = 20*, *minmapqmax = 15*, and *minstrandratio = 0.1*. No other variant filters were applied except for *rarity* and *minallelefraction* parameters, which were lowered compared to Illumina data (0.45 instead of 0.51) to account for the higher error rate of the Nanopore reads. Finally, as including a pairing rate in score calculation was not applicable, *usepairing* was set to *false*. The resulting .vcf files were processed and the consensus sequences generated as described for Illumina data.

#### 2.7.3. *De novo* assembly of Nanopore data

Fully processed Nanopore reads were used for *de novo* assembly with Canu v.1.8 [48]. The program was run with the following parameters: the estimated genome size was set to 180 kilobases (kb), somewhat above the 151 kb known from *A. frigida*. To account for the small overlaps present between some of the long-range PCR fragments, *minOverlapLength* was lowered to 300 bp. Correspondingly, the minimum length for reads being available for assembly was set to 300 bp as well. The *correctedErrorRate* was lowered to 0.134 and the three *MhapSensitivity* parameters set to *low*, as recommended by the program’s parameter reference for datasets with high coverage > 60-fold. *corOutCoverage* was set to 6,000, i.e. higher than the total input coverage to ensure that in principle, all loaded reads would be corrected. However, to avoid a detrimentally high coverage of reads during assembly (see Discussion, section 4.6.) while assuring that the program uses the longest reads first, *readSamplingCoverage* was given a value of 400 and *readSamplingBias* was set to 2.0, which results in the shortest reads (initially loaded into the sequence store) being flagged and excluded from analysis until a coverage of 400-fold remains. After assembly, resulting contigs supported by fewer than 10 reads and / or with a read coverage below 5-fold for more than 50% of their length were filtered by Canu. For checking purposes, *stopOnReadQuality = true* was applied. The remaining settings were kept on default. Assembled contigs were first locally blasted (megaBLAST) against the *A. frigida* reference without the IR_A_ to assure that no chimeric or contaminant contigs were present. Chimeric contigs with all parts mapping to *A. frigida* were split with the *fastasubseq* option in Exonerate [49] and (where necessary) reverse-complemented for further use. Contigs were then polished with Nanopolish v.0.10.2 (available at https://github.com/jts/nanopolish; [11]) as follows, using the original .fast5 files from the sequencing run. First, Minimap2 v.2.14 [50] with setting *-ax map-ont* was used for mapping raw reads to assembled contigs. After indexing the mapping with SAMtools, the *nanopolish variants* command was executed on each contig in consensus-calling mode, with *--max-haplotypes* set to 30,000 and *--min-candidate-frequency* to 0.2. Based on the extracted variants, the improved contigs were then generated via *nanopolish vcf2fasta*. Polished contigs were mapped to the *A. frigida* IR_A_-free reference with Minimap2, using setting *--ax asm20*, and their correct order and starting position identified in IGV. Merging of the contigs, together with the Sanger sequence of the gap between long-range PCR fragments 14 and 15 was done in BioEdit v.7.2.5 [51]. In one case, where the overlapping regions between two contigs were not identical but contained five single basepair differences, one of the contigs was arbitrarily chosen over the other. As the resulting assembly still lacked the IR_A_, the latter was added by copying and reverse-complementing the respective sequence of the IR_B_ and pasting it at the end of the assembled sequence after the SSC. Read coverage values were obtained by mapping the reads used by Canu for the first assembly step after correction and trimming (the *trimmedReads.fasta*) to the *de novo* assembled cp genome sequence without the IR_A_, using the *mapPacBio.sh* script as described in section 2.7.2. but setting *maxindel = 100*. Additionally, Qualimap was used for plotting coverage across the reference.

#### 2.7.4. Hybrid *de novo* assembly of Nanopore and Illumina data

Hybrid *de novo* assembly was performed using two approaches, once by analyzing reads from both platforms simultaneously in Unicycler, which also supports hybrid assembly, and once by using the information contained in high-quality Illumina data for improving the error-rich Nanopore raw reads before assembly. For this, fully processed Nanopore reads from *L. vulgare* were corrected based on sequence information from corresponding processed short Illumina reads using Nanocorr v.0.01 [52]. The resulting reads were used for an improved *de novo* assembly with Canu. Settings were kept identical to the assembly with unimproved reads, with a few exceptions. Due to their now increased quality, higher read coverage is not expected to be as problematic as with unimproved reads; hence, coverage was only limited by setting *minReadLength* to 2000; *readSamplingBias* was turned off by setting it to 0.0 (meaning that *readSamplingCoverage* being set to 200 would have no effect). Additionally, for these settings *stopOnReadQuality* was set to *false*. Assembled contigs were processed and merged and the final assembled sequence generated as described in section 2.7.3. However, no polishing was performed since polishing with the original, unimproved reads could have deteriorated the quality of the contigs.

### 2.8. Annotation of the chloroplast genomes

The two complete, finished sequences from Illumina *de novo* assemblies of *L. vulgare* and *L. virgatum* were annotated using the online tool GeSeq [53] available at https://chlorobox.mpimp-golm.mpg.de/geseq.html. The BLAT [54] searches were done setting the protein search identity to 80% and the rRNA, tRNA and DNA search identity to 85%. As references, two closely related chloroplast genomes (from tribe Anthemideae) available in Genbank were used: *A. frigida* and *Soliva sessilis* Ruiz & Pav. (subtribe Cotulinae; Genbank ID NC_034851.1). For chloroplast CDS and rRNAs, the curated MPI-MP Embryophyta chloroplast references were used as well. A HMMER profile search and tRNA annotation using ARAGORN v.1.2.38 [55] (with standard settings as on the website) were conducted additionally to check for possible annotation conflicts. After automatic annotation, putative annotation errors or discrepancies resulting from different references or algorithms were solved manually, deciding in favor of the closest references or the majority of algorithms / (published) references. The .gb file was also edited to include the very small 5’-exons of the *petB*, *petD* and *rpl16* genes. The final annotated chloroplast genomes were converted into graphical maps using the OGDRAW [56] online tool [57], available at https://chlorobox.mpimp-golm.mpg.de/OGDraw.html, with standard settings except that introns were also included in the genome map.

### 2.9. Comparison of *L. vulgare* assembly results from different methods to the Illumina *de novo* assembly

Finalized *L. vulgare* chloroplast genome sequences resulting from the different sequencing and analysis pipelines (Illumina, Nanopore, mapping assembly and *de novo* assembly) were compared to the Illumina *de novo* assembly. The same was done for the *L. virgatum* Illumina *de novo* assembly and the *Artemisia frigida* reference. All comparisons were calculated without the IR_A_. Each pair of sequences was first aligned online using MAFFT v.7 [58, 59] (available at https://mafft.cbrc.jp/alignment/server/), employing the FFT-NS-2 progressive method and otherwise standard settings. The alignment was adjusted to the length of the Illumina *de novo* reference, either by deleting bases from or by adding N’s to the sequence to be compared. The prepared alignments were then analyzed using the shell script *alignment_info2.sh* (written by Ulrich Lautenschlager, Regensburg, available upon request), which identifies the total number of mismatches, substitutions, insertions and deletions between two sequences. The length and G+C content of each assembly was identified using BioEdit.

### 2.10. Sequence divergence and potential molecular marker identification

Apart from comparison with *alignment_info2.sh* as outlined above, sequence variation between the *L. vulgare* and *L. virgatum* Illumina *de novo* assemblies was assessed via mapping processed *L. virgatum* Illumina reads to the *L. vulgare* Illumina *de novo* assembly. Mapping, read deduplication and variant calling was done according to section 2.6.2.; during mapping, *maxindel* was set to 300 in BBMap. After packing and indexing, the resulting .vcf file was analyzed regarding variant density with VCFtools. The number of variants was output in 1,000 bp windows and graphically illustrated. Based on the highest peaks in the resulting graph (windows containing more than eight variants), the most variable regions in the cp genome were identified and the corresponding molecular markers (intergenic spacers / genes) examined regarding total variant numbers and variation in relation to marker length.

## 3. Results

### 3.1. Long-range PCR, sequencing, and genome assembly

The lengths of long-range PCR fragments obtained from *Leucanthemum vulgare* and *L. virgatum* were similar to those found in *Artemisia frigida*, with the shortest and longest fragments in *L. vulgare* being 6,086 bp (fragment 4) and 12,499 bp (fragment 14) long, respectively. Read statistics for all three runs are shown in Table 1. Sequencing of *L. virgatum* resulted in a total of 381,932 raw reads. 0.7% contaminant reads were removed, and length and quality filtering reduced the dataset by another 43.8%, leaving 212,186 trimmed and filtered high-quality reads (55.6%) for subsequent analyses. Sequencing of *L. vulgare* yielded 373,518 raw reads, of which 1.5% belonged to the PhiX spike-in and 32% were removed during length and quality filtering. Thus, 248,456 reads (66.5%) remained for analysis. Approximately 1.5 hrs sequencing of *L. vulgare* on a flow cell with two multiplexed samples using Oxford Nanopore MinION resulted in a total yield of 632.9 megabases (Mb) with a mean read quality of 7.49. A fraction of 22.6% from the raw reads within the Albacore “pass” folder were discarded by Porechop due to middle adapters. After demultiplexing and adapter trimming by Porechop, 52,223 *L. vulgare* reads with a total of 161,521,432 bp were obtained, with read lengths ranging from 101-26,011 bp, an average read length of 3,093 bp and mean sequence qualities ranging from Q8 - Q16 (with 48.8% of the reads having Q12 or Q13). A total of 217 reads were quality- and length-filtered, leaving 52,006 reads for mapping assemblies. For *de novo* assembly, 19,111 putative contaminant reads were additionally removed, leaving 32,895 reads. Chimera-splitting by Pacasus raised that number to 59,874 reads, of which 19,311 reads had lengths below 50 bp and were excluded. Thus, 40,563 reads were available for *de novo* assembly; these had an average read length lowered by about 750 bp compared to the contaminant-removed reads. For the hybrid *de novo* assembly, read improvement by Nanocorr lowered the read number to 38,107, but added a total of 675,084 bp to the remaining reads. All sequenced raw reads which were analyzed in the course of the present study are available at the NCBI Sequence Read Archive (SRA) under accession numbers XX, XX and XX.

**Table 1.** Read statistics for Illumina and Nanopore sequencing runs.

|  | *L. virgatum*<br>Illumina | *L. vulgare*<br>Illumina | *L. vulgare*<br>Nanopore |
| --- | --- | --- | --- |
| raw reads (300+300 run / 80+80 run) | 381,932<br>(25,290 / 356,642) | 373,518<br>(22,810 / 350,708) | 52,223 |
| contaminant reads removed (%) | 2,620 (0.7) | 5,560 (1.5) | 0 (0.0) |
| quality- and length-filtered reads (%) | 167,126 (43.8) | 119,502 (32.0) | 217 (0.4) |
| remaining reads for analysis (%) | 212,186 (55.6) | 248,456 (66.5) | 52,006 (99.6) |

Number of raw reads is given for each of both MiSeq 300-bp and 80-bp paired-end sequencing runs. Length-filtered reads include those which were too short after trimming. Read numbers for Nanopore contaminant and filtered reads as well as remaining reads include only processed, not fully processed reads (see text).
*L.*, *Leucanthemum*.

For the mapping assemblies, detailed statistics are available in Table 2. Mapping Nanopore data to the *Artemisia* reference yielded one multiallelic site in the .vcf file, which was resolved manually by discarding the variant with the lower quality in the file. All other observed variants were biallelic. Mapped reads comprised 95.7-98.2% of the input data, with the percentage of ambiguously mapped reads being higher in Illumina (14.6% on average) than in Nanopore reads (5.97%). Mean coverage was well over 100-fold in all mapping assemblies, but with large standard deviations. However, over 99% of the reference still had coverage ≥ 30-fold for Illumina and ≥ 100-fold for Nanopore data. Variant calling resulted in 379 variants with a mean depth of 102.42-fold and a mean variant quality of 50.7 for the Illumina mapping (*virgatum*), 1,925 variants (depth 112.98-fold, variant quality 50.66) for the Illumina mapping (*Artemisia*), 297 variants (depth 685.48-fold, variant quality 16.77) for the Nanopore mapping (*virgatum*) and 1,645 variants (depth 721.59-fold, variant quality 16.72) for the Nanopore mapping (*Artemisia*).

**Table 2.** Mapping statistics for reference-based assembly of Illumina and Nanopore data.

|  | Illumina<br>(*L. virgatum*) | Illumina<br>(*Artemisia*) | Nanopore<br>(*L. virgatum*) | Nanopore<br>(*Artemisia*) |
| --- | --- | --- | --- | --- |
| input reads | 248,456 | 248,456 | 39,080 | 39,080 |
| mapped reads (%) | 238,574 (96.02) | 237,686 (95.67) | 38,377 (98.20) | 38,363 (98.17) |
| ambiguously mapped (%) | 36,641 (14.75) | 35,934 (14.46) | 2,341 (5.99) | 2,325 (5.95) |
| read lengths | 50-300 bp | 50-300 bp | 1-3,000 bp | 1-3,000 bp |
| mean coverage (stdev) | 143.41x<br>(88.36) | 143.43x<br>(87.92) | 754.34x<br>(473.68) | 756.68x<br>(472.80) |
| fraction of reference at<br>coverage x or higher | 30x: 99.74%<br>20x: 99.88% | 30x: 99.66%<br>20x: 99.81% | 100x: 99.72%<br>50x: 99.76% | 100x: 99.72%<br>50x: 99.76% |
| mean mapping quality | 36.5 | 33.4 | 22.4 | 19.6 |
| duplicate reads removed | 7,644 | 7,646 | n.a. | n.a. |
| remaining reads after<br>deduplication (%) | 230,930 (96.8) | 230,040 (96.8) | n.a. | n.a. |

The respective reference used for mapping (*L. virgatum* Illumina *de novo* assembly or *A. frigida* as obtained from Genbank, both without the IR<sub>A</sub>) is given in parentheses. Mapped reads include ambiguously mapped reads. The percentages given for both refer to all input reads, the fraction of remaining reads after deduplication is relative to mapped reads. No deduplication was performed for Nanopore reads. Nanopore read lengths range from 1-3,000 bp due to necessary splitting of long reads into 3,000-bp chunks by the mapping software.
*L.*, *Leucanthemum*; stdev, standard deviation; n.a., not applicable.

*De novo* assembly of Illumina data from *L. virgatum* produced five contigs (see Table 3), of which three belonged to the IR; one unnecessary connection between two IR contigs had to be removed before doubling and merging. *Leucanthemum vulgare* data yielded four contigs, one of which (a small contig of 1,916 bp, see Discussion section 4.6.) did not blast as chloroplast sequence and therefore was excluded. The remaining three fragments corresponded to the LSC, SSC, and IR. Fragment lengths in both species varied between 621 and 82,675 bp. Mean read coverages were 128.30-fold and 152.28-fold, respectively. In *L. vulgare*, only 0.26% of the assembled sequence was covered at below 30-fold while in *L. virgatum*, 6.62% of the sequence length had lower coverage. However, 98.66% of the assembly still was covered at least 20-fold. *De novo* assembly of Nanopore-only data in *L. vulgare* started with 8,338 reads for correction and used 4,632 reads for the actual assembly, which resulted in six contigs. One of those was discarded as it was a chimera with one of two parts being a contaminant which did not blast to *Artemisia* at all (see Discussion section 4.6.); another was split into two parts, resulting in six final contigs with fragment lengths between 6,655 and 53,891 bp. Polishing rewrote the contigs with 118 substitutions, 1,100 insertions and 46 deletions. Contigs were merged together with the 480-bp Sanger sequence of the primer gap (see section 2.5.). For the hybrid *de novo* assembly, 14,351 reads were used for correction and 10,422 reads for assembly; the latter yielded five contigs, of which two had to be split. The seven final contigs had lengths between 6,819 and 54,677 bp. Mean read coverage was considerably higher for the Nanopore-only assembly (321.43-fold) than for the Illumina assemblies; nevertheless, 8.83% of the assembled sequence was covered by 99 or fewer bases. However, coverage was at least 50-fold for 99.83% of the assembly. For the hybrid assembly, where there was less need for keeping coverage low (559.39-fold), the amount of bases at coverage below 100-fold accordingly dropped to 0.29%.

**Table 3.** De novo assembly statistics.

|  | *L. virgatum*<br>Illumina | *L. vulgare* Illumina | *L. vulgare*<br>hybrid assembly | *L. vulgare*<br>Nanopore |
| --- | --- | --- | --- | --- |
| input reads | 212,186 | 248,456 | 10,422 | 4,632 |
| best k-mer | 65 | 55 | n.a. | n.a. |
| no. of *Leucanthemum*<br>contigs (after splitting) | 5 | 3 | 5 (7) | 5 (6) |
| contig sizes | 621-82,641 bp | 18,400-82,675 bp | 6,819-54,677 bp | 6,655-53,891 bp |
| mean coverage (stdev) | 128.30x<br>(109.88) | 152.28x<br>(113.00) | 559.39x<br>(323.29) | 321.43x<br>(167.74) |
| fraction at coverage x<br>or higher | 30x: 93.38%<br>20x: 98.66% | 30x: 99.74%<br>20x: 99.91% | 100x: 99.71%<br>50x: 99.77% | 100x: 91.17%<br>50x: 99.83% |

Three *de novo* assemblies were produced for *L. vulgare*, using Illumina data, Nanopore data or Nanopore data improved by Illumina data (hybrid approach). Assembly was done with Unicycler for Illumina reads and Canu for Nanopore reads. Nanopore input reads are given after Canu correction and trimming steps. Values for 'fraction at coverage x or higher' refer to the final assembled sequence without the IR<sub>A</sub> and denote the percentage with a certain read coverage after mapping of input reads.
*L.*, *Leucanthemum*; stdev, standard deviation; no, number; n.a., not applicable.

### 3.2. Genome organization, features and gene content

The *Leucanthemum vulgare* chloroplast genome (Fig 2) as assembled *de novo* from Illumina reads is 150,191 bp in length, the *L. virgatum* plastome is 150,120 bp long (Supporting Information S3 Fig); their overall G+C content is 37.45% (Supporting Information S4 Table). The genome sequences alongside their annotations have been deposited in NCBI’s Genbank under accession numbers XXXXX and XXXXX. The two cp genomes are highly similar, featuring an LSC region of 82,675 and 82,641 bp, respectively, an SSC region of 18,400 vs. 18,435 bp, and a pair of inverted repeats of each 24,558 bp in *L. vulgare* and 24,522 bp in *L. virgatum* (Supporting Information S4 Table). The LSC/IR/SSC boundaries of both species as determined by Sanger sequencing are depicted in Fig 3. In *L. vulgare*, the IR extends slightly further into the *rps19* and *ycf1* genes compared to *L. virgatum*. In the latter, the SSC region was assembled in reverse-complement direction to *L. vulgare* (see Discussion section). Both genomes possess the two inversions typical for Asteraceae: one large (Inv 1) and one small inversion (Inv 2) with respect to *Nicotiana tabacum* L. and other non-Asteroid families, located within the LSC [31, 32].

**Fig 2.**
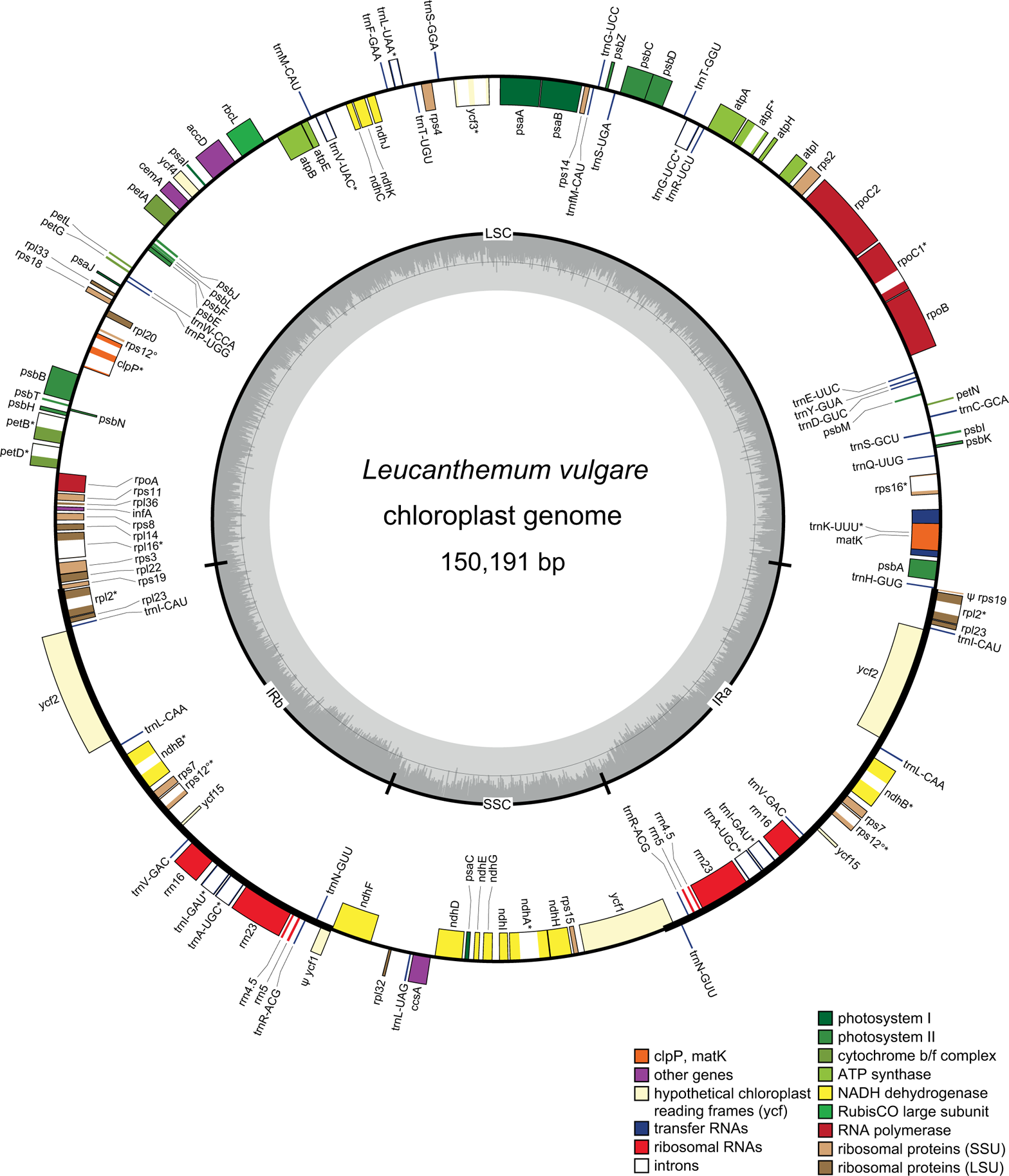
Chloroplast genome map for *Leucanthemum vulgare*. Genes on the outside of the outer circle are transcribed counterclockwise, genes on the inside are transcribed clockwise. Introns are illustrated with white color within genes; genes containing an intron are additionally marked with *. Pseudogenes are preceded by a ψ. The trans-spliced *rps12* gene is marked with °. Color-coding of genes depicts their affiliation to the functional groups given. The inner circle indicates the borders of the large single-copy (LSC) and small single-copy (SSC) regions as well as the inverted repeats (IR). The innermost gray shaded area shows the G+C content of the cp genome. The gene order is identical in *L. virgatum* (see Supporting Information S3 Fig), whereas their exact positions and the extent of the inverted repeat slightly differ (Fig 3).

**Fig 3.**
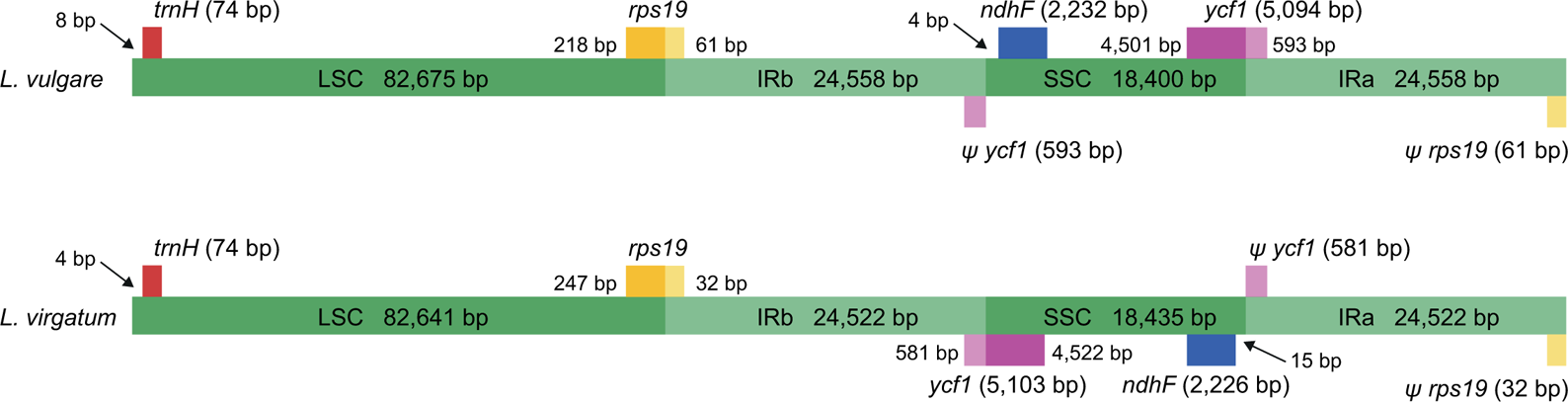
Comparison of the inverted repeat (IR) boundaries in *Leucanthemum vulgare* and *L. virgatum*. Genes above the green bars are transcribed in reverse direction, those below in forward direction. For both taxa, the IR extends into the *ycf1* and *rps19* genes, resulting in two pseudogenes (denoted by a ψ) at the single-copy / IR junctions. Lengths are given for whole genes and their duplicated fragments as well as the large single-copy (LSC), small single-copy (SSC) and IR regions (IRA and IRB). Arrows show basepair (bp) distance from the junctions for the *ndhF* and *trnH* genes. The SSC in *L. virgatum* is exemplarily reverse-complemented with respect to that in *L. vulgare*; both configurations exist in individual plants according to [73]. The figure is not to scale.

The arrangement and order of genes is identical in the two species: the plastomes contain 80 predicted protein-coding genes, four rRNA genes, and 30 tRNA genes coding for all 20 amino acids. Seven protein-coding genes, all rRNA genes and seven tRNA genes are duplicated due to the IR, raising the total number of genes to 132. Six of the 30 tRNA genes and 12 of the 80 protein-coding genes contain introns; of the latter, two harbor two introns while the rest has one single intron. Two pseudogenes are present in *Leucanthemum*: ψ-*ycf1* and ψ-*rps19* are located at the IR/SC boundaries (Fig 3), their lack of functionality being due to only partial duplication. The trans-spliced *rps12* gene is found in the LSC (5’-end) and the IR regions (duplicated 3’-end). A summary of all genes is given in Table 4.

**Table 4.**
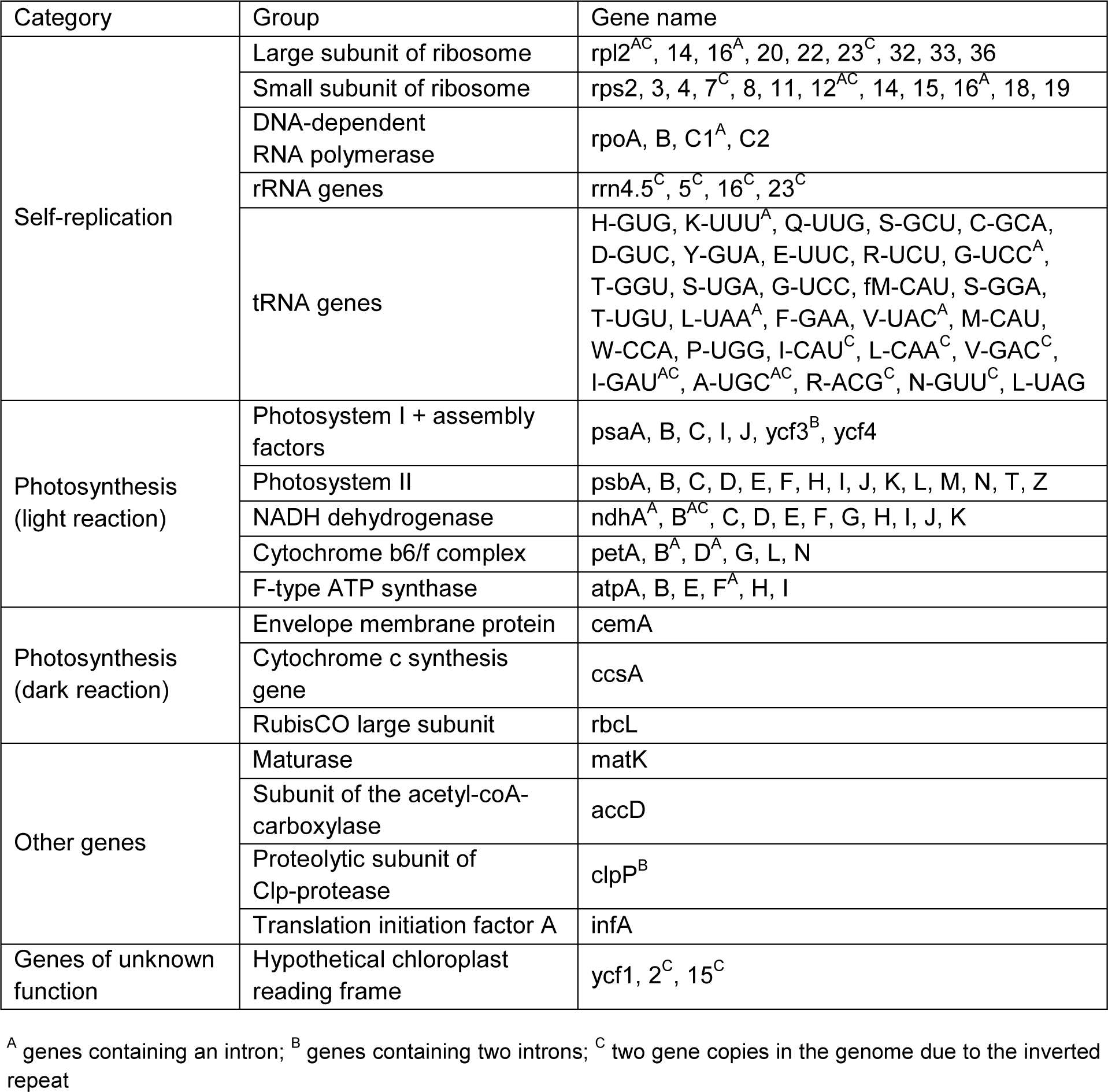
Genes present in the sequenced Leucanthemum vulgare and L. virgatum genomes.

| Category | Group | Gene name |
| --- | --- | --- |
| Self-replication | Large subunit of ribosome | rpl2<sup>AC</sup>, 14, 16<sup>A</sup>, 20, 22, 23<sup>C</sup>, 32, 33, 36 |
|  | Small subunit of ribosome | rps2, 3, 4, 7<sup>C</sup>, 8, 11, 12<sup>AC</sup>, 14, 15, 16<sup>A</sup>, 18, 19 |
|  | DNA-dependent RNA polymerase | rpoA, B, C1<sup>A</sup>, C2 |
|  | rRNA genes | rrn4.5<sup>C</sup>, 5<sup>C</sup>, 16<sup>C</sup>, 23<sup>C</sup> |
|  | tRNA genes | H-GUG, K-UUU<sup>A</sup>, Q-UUG, S-GCU, C-GCA, D-GUC, Y-GUA, E-UUC, R-UCU, G-UCC<sup>A</sup>, T-GGU, S-UGA, G-UCC, fM-CAU, S-GGA, T-UGU, L-UAA<sup>A</sup>, F-GAA, V-UAC<sup>A</sup>, M-CAU, W-CCA, P-UGG, I-CAU<sup>C</sup>, L-CAA<sup>C</sup>, V-GAC<sup>C</sup>, I-GAU<sup>AC</sup>, A-UGC<sup>AC</sup>, R-ACG<sup>C</sup>, N-GUU<sup>C</sup>, L-UAG |
| Photosynthesis (light reaction) | Photosystem I + assembly factors | psaA, B, C, I, J, ycf3<sup>B</sup>, ycf4 |
|  | Photosystem II | psbA, B, C, D, E, F, H, I, J, K, L, M, N, T, Z |
|  | NADH dehydrogenase | ndhA<sup>A</sup>, B<sup>AC</sup>, C, D, E, F, G, H, I, J, K |
|  | Cytochrome b6/f complex | petA, B<sup>A</sup>, D<sup>A</sup>, G, L, N |
|  | F-type ATP synthase | atpA, B, E, F<sup>A</sup>, H, I |
| Photosynthesis (dark reaction) | Envelope membrane protein | cemA |
|  | Cytochrome c synthesis gene | ccsA |
|  | RubisCO large subunit | rbcL |
| Other genes | Maturase | matK |
|  | Subunit of the acetyl-coA-carboxylase | accD |
|  | Proteolytic subunit of Clp-protease | clpP<sup>B</sup> |
|  | Translation initiation factor A | infA |
| Genes of unknown function | Hypothetical chloroplast reading frame | ycf1, 2<sup>C</sup>, 15<sup>C</sup> |

<sup>A</sup> genes containing an intron; <sup>B</sup> genes containing two introns; <sup>C</sup> two gene copies in the genome due to the inverted repeat

### 3.3. L. vulgare and L. virgatum sequence divergence and molecular marker identification

The *L. virgatum* plastome has 99.17% nucleotide identity to *L. vulgare* (see Supporting Information S4 Table), the cp genome of *Artemisia frigida* is 96.31% identical. Alignment of both genomes yielded a total of 4,704 mismatches, while *L. virgatum* differs from *L. vulgare* in only 1,038 mismatches, 28.8% of which are substitutions, 37.7% are gaps, and 33.5% are inserted bases. Analysis of the variant density of *L. virgatum* with respect to *L. vulgare* showed six regions of high variance (Fig 4), corresponding to three intergenic spacers and two genes (*ndhF* and *ycf1*), which vary in 8-28 positions and have a variability of 0.55-1.38% (Table 5). The most promising intergenic spacer was *trnE-rpoB*, with 12 variants within an easily amplifiable length of 869 bp. The *ycf1* gene features two variability hotspots of 794 bp (1.39% variability) and 488 bp (1.84% variability), respectively; at the end of the *ndhF* gene, one hotspot of 750 bp with 1.6% variability was found.

**Fig 4.**
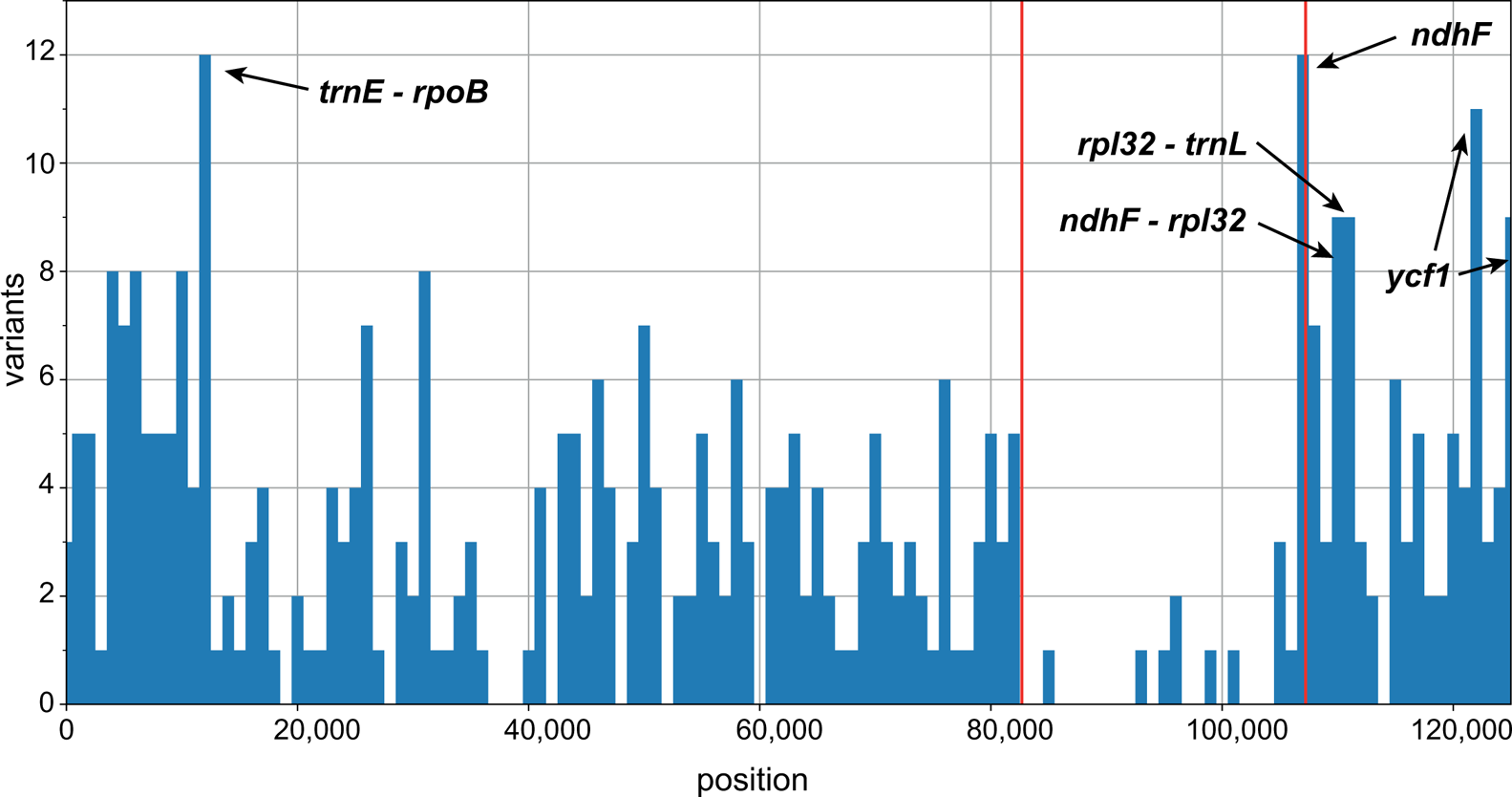
Nucleotide variant density in *L. vulgare* and *L. virgatum* chloroplast genomes. Illumina reads from *L. virgatum* were mapped to the Illumina *de novo* assembled sequence of *L. vulgare* (lacking the second inverted repeat) and variants called. Plot data on variant density generated by VCFtools in 1000-bp windows. The position of the first inverted repeat (IRB) is indicated by red lines. Peaks depicting potentially useful marker candidates for *Leucanthemum* are highlighted by arrows.

**Table 5.** Chloroplast genomic regions with the highest variability between *L. vulgare* and *L. virgatum*.

| Marker | Position [bp] | Length [bp] | Variants | Variation [%] |
| --- | --- | --- | --- | --- |
| *trnE<sup>UUC</sup> – rpoB* | 11,909 – 12,778 | 869 | 12 | 1.38 |
| *ndhF*<br>*ndhF* hotspot region 1 | 107,238 – 109,469<br>107,249 – 107,999 | 2,231<br>750 | 19<br>12 | 0.85<br>1.60 |
| *ndhF – rpl32* | 109,470 – 110,488 | 1,018 | 8 | 0.79 |
| *rpl32 – trnL<sup>UAG</sup>* | 110,654 – 111,640 | 986 | 11 | 1.12 |
| *ycf1*<br>*ycf1* hotspot region 1<br>*ycf1* hotspot region 2 | 121,133 – 126,226<br>122,119 – 122,913<br>125,081 – 125,569 | 5,093<br>794<br>488 | 28<br>11<br>9 | 0.55<br>1.39<br>1.84 |

### 3.4. *De novo* and mapping assemblies of *L. vulgare* with Illumina and Nanopore data

The seven assembly methods tested on *L. vulgare* resulted in plastome sequences of lengths ranging between 125,633 bp (100%) in the Illumina *de novo* assembly and 124,851 bp in the Illumina mapping (*Artemisia*) assembly (Table 6). The sequences (except from the Illumina *de novo* assembly which was deposited in Genbank, see above) are available in Supporting Information S5 File. Setting Illumina *de novo* as the gold standard, the Nanopore hybrid assembly achieved the highest identity with the former (99.98%). Most divergent from the standard was the Nanopore mapping (*Artemisia*) with 98.29%. The quality of the Nanopore mapping approach increased when using the more closely related reference *L. virgatum* (99.51%), which is almost equal to the Nanopore *de novo* assembly (99.59%) and better than mapping the high-quality Illumina data to the more distantly related *Artemisia* (99.27%). Generally, *de novo* assembly always yielded better results than mapping assembly. Unlike in the comparison between the two *Leucanthemum* species, where substitutions, gaps, and inserted bases contribute more or less equally to the total number of mismatches (Supporting Information S4 Table), deletions are the main source of error in all tested approaches, accounting for 56.2-96.0% of the differences. Nanopore mapping seemed to yield the highest proportion of insertion errors (32.1-34.6%), while, compared to the other approaches, most erroneous substitutions were introduced by Nanopore *de novo* assembly (18.5%).

**Table 6.** Comparison of six assemblies of the Leucanthemum vulgare plastome, obtained using different methods and data types, with a “gold-standard” Illumina de novo assembly.

|  | Illumina<br>*d.n.* | Illumina<br>map (virg) | Illumina<br>map (Art) | Nanopore<br>hybrid *d.n.* | Nanopore<br>*d.n.* | Nanopore<br>map (virg) | Nanopore<br>map (Art) |
| --- | --- | --- | --- | --- | --- | --- | --- |
| assembly length<br>without IR<sub>A</sub> [bp] | 125,633 | 125,383 | 124,851 | 125,609 | 125,270 | 125,451 | 125,165 |
| % GC content | 36.35 | 36.38 | 36.45 | 36.36 | 36.37 | 36.36 | 36.43 |
| % identity to<br>*L. vulgare de novo* | n.a. | 99.74 | 99.27 | 99.98 | 99.59 | 99.51 | 98.29 |
| % alignment<br>positions with N | n.a. | 0 | 0 | 0 | 0.12 | 0 | 0 |
| total mismatches | n.a. | 329 | 922 | 25 | 362 | 619 | 2,164 |
| no. of substitutions<br>(% total mismatches) | n.a. | 19 (5.8) | 56 (6.1) | 1 (4.0) | 67 (18.5) | 39 (6.3) | 198 (9.1) |
| no. of gaps<br>(% total mismatches) | n.a. | 280 (85.1) | 824 (89.4) | 24 (96.0) | 253 (69.9) | 381 (61.6) | 1217 (56.2) |
| deletion events | n.a. | 21 | 47 | 5 | 221 | 57 | 149 |
| no. of inserted bases<br>(% total mismatches) | n.a. | 30 (9.1) | 42 (4.6) | 0 (0.0) | 42 (11.6) | 199 (32.1) | 749 (34.6) |
| insertion events | n.a. | 5 | 3 | 0 | 42 | 42 | 67 |

Two data types (Illumina, Nanopore reads) and three analysis pipelines were used to assemble the *L. vulgare* plastome: *de novo* assembly, reference-based (mapping) assembly and hybrid *de novo* assembly using Nanopore reads corrected by Illumina data. Comparisons were made based on alignment of the respective assembly to *L. vulgare de novo* (Illumina). Percentages of identity and 'alignment positions with N' are referable to the length of the alignment; 'alignment positions with N' denotes the amount of Ns required for alignment to the Illumina *de novo* assembly.
*L.*, *Leucanthemum*; IR<sub>A</sub>, inverted repeat A; n.a., not applicable; *d.n.*, *de novo*; map, mapping assembly; virg, *L. virgatum* Illumina *de novo* assembly used as reference for mapping; Art, *Artemisia frigida* as obtained from Genbank used for mapping; both references without the IR<sub>A</sub>.

## 4. Discussion

### 4.1. Enrichment of chloroplast DNA using long-range PCR

For the present study, we relied on enrichment of cp DNA via long-range PCR. For this approach, several primer sets are available (e.g., [60], producing nine fragments). The protocol chosen here [28] resulted in 16 fragments with a maximum length of 12,499 bp. Compared with other approaches, the advantage of long-range PCR is that in most cases, it will exclusively enrich for plastid DNA, including segments possibly transferred there from the nucleus or mitochondrion, while omitting nuclear inserts of plastid DNA and requiring substantially less leaf material. Of course, the inclusion of paralogous cp genes from the nucleus or mitochondrion cannot be fully ruled out [2]. Another limitation is that highly degraded DNA cannot be used for long-range PCR, which limits its use with herbarium material [61, 6]. The method is also unsuitable if larger rearrangements in gene order are to be expected, as have been found for example in *Passiflora* L. [62]. In any case, optimization of primers for the study group will almost always be necessary to amplify all fragments (in the present study, different primers had to be used for five out of 16 fragments, and several primers were slightly modified). It is also essential to use a high-quality-, proofreading polymerase to avoid misinterpretation of erroneously incorporated bases as true variants, and to remove unwanted primer sequence during read processing.

The main disadvantage of PCR-based enrichment however seems to be the resulting uneven read coverage, which is partly due to doubled sequencing at PCR-fragment overlaps. However, apart from that, although fragments were pooled at equimolar ratios before sequencing in the present study, fragment coverage across the cp genome turned out to be very variable for Illumina as well as Nanopore reads (Fig 5). Possible reasons for such a behavior include adapter ligation biases and differences among targets regarding denaturation and annealing during library preparation [2]. The same effect was mentioned by [63], who by contrast found much lower variation in coverage when using DNA from isolated chloroplasts. Altogether, long-range PCR thus cannot be regarded the optimal approach for plastome sequencing, also because exceptionally high coverage in only a subset of regions might cause problems during Nanopore *de novo* assembly (see section 4.6.). A suitable alternative might be “genome skimming” [64, 6] or solution-based hybridization (“target enrichment”) via custom-designed oligonucleotide probes representing the cp genome [65, 66]. However, both approaches require a closely related reference for chloroplast read extraction or bait design. Chloroplast DNA can also be obtained by isolating whole chloroplasts before sequencing. However, a high amount of fresh leaves is required for isolation [6], which may exceed 5 g or even 20 g [67].

**Fig 5.**
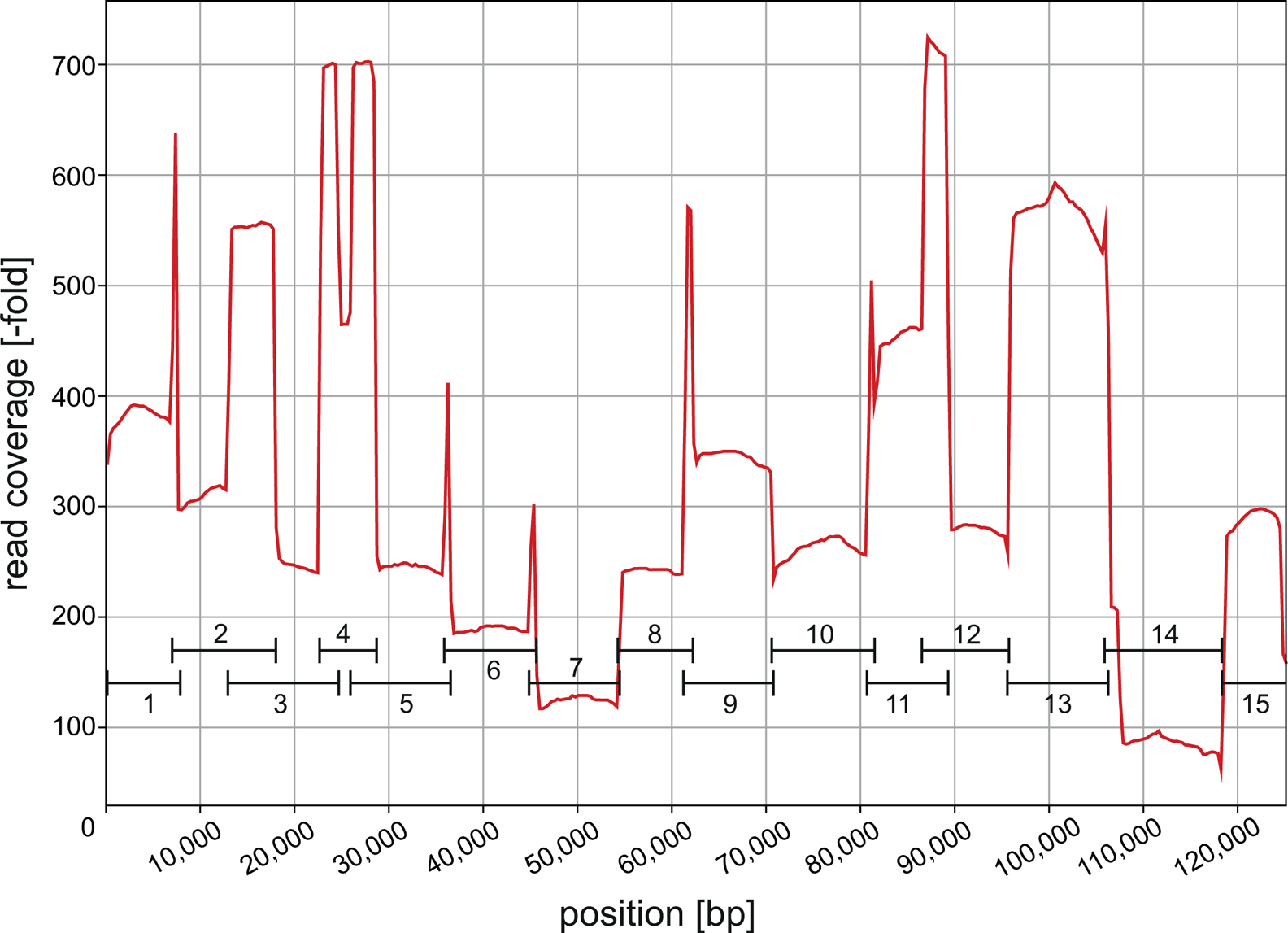
Chloroplast genome read coverage of the Nanopore *de novo* assembly. *Leucanthemum vulgare* Nanopore long reads used for assembly (post-correction and post-trimming) were mapped to the assembled sequence lacking the second inverted repeat (IR_A_) and coverage-across-reference plot data extracted with Qualimap. Black lines represent the long-range PCR fragments reads are based on. Note the uneven coverage across fragments and coverage peaks at fragment overlaps. bp, basepairs.

### 4.2. Assembly of chloroplast genomes using NGS

The assembly of cp genomes using Illumina short reads is still regarded as the “gold standard” regarding data quality and integrity. The most common outcome of *de novo* plastid assembly today are three long contigs corresponding to the LSC, SSC, and IR regions [6], as presented for example in [21] and the present study. The G+C content of the two Illumina *de novo* assemblies presented here is typical for cp genomes [68] and suggests that no large portions of nuclear DNA have been erroneously incorporated. Both plastomes are highly similar (0.83% sequence divergence); as either species belongs to one of two main clades found in *Leucanthemum* [69, 70], this might represent the average cp genome divergence within the genus. The cp genome of *Artemisia frigida*, belonging to a different subtribe, still shows only 3.69% divergence from *L. vulgare*, which points toward close relationships within Anthemideae as a whole. The SSC region of *L. virgatum* was assembled to be reverse-complemented with regard to that in *L. vulgare*. While such differences have been highlighted as “unique sequence rearrangement event” by [71] and also in other studies (see [72]), it has already been shown by [73] that molecules of both SSC orientations exist equimolarily within the same individual. It is also important to realize that with the use of PCR fragments that do not span the entire IR, it is *per se* impossible to distinguish between the two orientations.

The comparison of the two new plastomes with regard to sequence variability yielded five potential markers. The intergenic spacers *trnE-rpoB*, *ndhF-rpl32* and *rpl32-trnL* were already described by other studies as being phylogenetically informative within angiosperms or specifically within Asteraceae [74, 75]. The *ycf1* gene has been recognized as an interesting region for barcoding, since it contains a high percentage of parsimony-informative characters, which are mostly due to the frequent occurrence of SSRs in the region [76, 77]. Another interesting aspect is the high variability at the end of the *ndhF* gene, in the region close to the IR/SSC boundary. However, sequencing of the entire *ycf1* or *ndhF* gene will not be too effective in terms of variability per sequenced length, so if their use as a molecular marker is intended, it would be advisable to design suitable primers around their hotspot regions. The generally lower substitution rate of the IR region, which has been demonstrated in several plant groups (see [78]), is attributed to its duplicative nature in combination with the effects of biased gene conversion [79, 80].

### 4.3. Hybrid assembly approaches

Three basic procedures can be used for hybrid genome assembly: either, an assembly is first generated based on Nanopore data alone and is subsequently polished with Illumina data (e.g., in [81]), often by using the tool Pilon. Alternatively, the recently published pipeline Unicycler follows a short-read-first approach, by assembling short reads to an assembly graph which is subsequently resolved using long reads. Third, long reads can be corrected by aligning short reads prior to assembly, for the benefit of providing high-quality input to the latter; this can be done by using the tool Nanocorr.

In the comparison of hybrid and non-hybrid approaches presented by Wang et al. (2018), the authors found that the hybrid approach performed best, yielding a single high-quality (final error rate: 0.0007 per base of a set of validation reads) contig representing the whole plastome, as long as 20-fold coverage of long and short reads was provided [21]. Hybrid assembly was also superior to Nanopore-only assembly in [82], where the authors sequenced a mitogenome from *Chrysanthemum* L. The same effect was observed in the present study. Hybrid assembly using Unicycler (results not shown) yielded results identical to *de novo* assembly of short reads alone (three contigs), but was not able to assemble a continuous contig covering the two IRs. This came as no surprise as the longest Nanopore read was too short with only 12,532 bp long, matching the length of the longest PCR fragment sequenced. The use of long-range PCR thus limits the possibilities of Unicycler and Nanopore reads in general. [21] also note that the performance of the hybrid assembly was “highly dependent on the length of the long-reads”: assembly of a single contig was already possible with a coverage of 5-fold of Nanopore reads of lengths > 30 kb (together with at least 8-fold short-read coverage), but failed when Nanopore read lengths were < 20 kb long (i.e., shorter than the IR region). Another hybrid approach pursued in the present study (correction of long reads using Nanocorr before assembly) resulted in five contigs of which two had to be split, and the final sequence was not completely identical to the Illumina *de novo* (and thus, the Unicycler hybrid) assembly (99.98%). This means that assembly based on short reads but supplemented with information from long reads produced better results than proceeding the other way around.

### 4.4. *De novo* assembly based on Nanopore data alone

Only very few studies have applied Nanopore sequencing exclusively for plastome assembly. Bethune et al. (2019) used Nanopore combined with target enrichment to sequence seven taxa from Poaceae and Palmaceae, of which one was subsequently used as reference for the others [24]. From the six assemblies which were conducted with the Flye assembler [83], none recovered the plastomes in a single contig, but two or more contigs of varying length. The authors [24] specifically noted the influence of input DNA quality on assemblies. DNA isolated from fresh tissue resulted in longer median read lengths and only two contigs with higher plastome coverage (with respect to the reference) than silica gel-dried samples. As mentioned above, this again underscores the importance of read length for Nanopore assembly. Wang et al. (2018), in their comparative study mentioned above, found that Canu was able to assemble the *Eucalyptus pauciflora* Sieber ex Spreng. plastome into one or two contigs depending on read coverage, which however lacked some genome regions or covered others twice [21]. No assembly covered the complete cp genome; the final error rate after polishing with Nanopolish decreased to 0.0022 (compared to 0.0007 in the hybrid assemblies). Our Nanopore-only assembly yielded slightly better results: the Canu assembly produced six contigs, which however covered all of the cp genome, except the first 152 bp. Although overlap among the contigs was up to ∼830 bp, no regions were covered twice. As a side note it is important to realize that despite the often advocated advantage of *de novo* assembly, namely no need for a reference genome, is not completely met for cases in which more than one contig is assembled. In our analyses, the six contigs all overlapped, but with very variable overlap lengths, with one overlap being as little as seven bp, five of which were a C-homopolymer. In such cases, reliable merging of contigs is often realized by mapping to a reference sequence (e.g., [84]; see also [85]), as was done here.

Although the Nanopore-only *de novo* assembly presented in our study could not fully keep up with that from the hybrid approach using Nanocorr (sequence identity to Illumina *de novo* 99.98%, see Table 6), it reached a comparatively high sequence identity of 99.59%. This is comparable to the final 99.5% nucleotide identity of the assembly by Loman et al. (2015) [11], whose methods were also applied in the present study (read correction with a POA graphs - dependent consensus algorithm as implemented in Canu, polishing of the final assembly with Nanopolish). Regarding the remaining errors in our assembly, almost 70% of all mismatches were gaps; the accumulation in deletion errors in homopolymer runs is clearly visible in visualizations of mappings prepared for the mapping approach and increases with homopolymer length. This fits the expectation for non-random Nanopore sequencing errors mentioned in [7].

### 4.5. Reference-based assembly using Nanopore data

Apart from *de novo* assembly, plastome sequences can also be obtained by simply aligning reads to a reference genome and creating a consensus sequence from the mapping. This approach is only suitable for studies of closely related taxa with uniform chloroplast genomes [2, 6]. The reason for that is that a mapping consensus will always adopt characteristics of the reference from which it was inferred. Structural rearrangements or additional transferred genes might easily be missed, especially when mapping short reads; the structure of repeat regions cannot be distinguished by mapping based approaches at all [86]. For the assembly of the globe artichoke (*Cynara* L.) cp genome with Illumina reads, [76] compared mapping and *de novo* approaches and found the results to be almost identical. [87] analyzed intraspecific diversity in *Brachypodium* P.Beauv. with Illumina data. According to their results, the reference-guided approach yielded fewer and longer contigs than *de novo* assemblies in most cases. In the present study, mapping assemblies based on Illumina data were found to be of somewhat lower quality, being 99.27% and 99.74% identical to the *de novo* assembly, respectively, depending on the mapping reference used.

However, to our knowledge, no study has yet employed Nanopore sequencing reads for reference-based assembly of plant chloroplast genomes. The mapping assemblies of *L. vulgare* presented here were produced using read lengths of up to 3,000 bp and two closely-related references, which do not show any structural reorganization compared to *L. vulgare* and also have the same gene order (nucleotide identities to *L. vulgare* Illumina *de novo*: *L. virgatum* 99.17%, *A. frigida* 96.31%). Illumina mappings were superior to Nanopore mappings for both references, but surprisingly, nucleotide divergence (from the Illumina *de novo* assembly) was only slightly worse in Nanopore mappings, with the difference between the two being as low as 0.23% when using the *L. virgatum* reference. However, this difference rose to 0.98% when using *Artemisia*. This loss in quality caused by using the more distantly related *Artemisia* instead of *L. virgatum* was found for both datatypes, but seemed to be worse with Nanopore: Illumina assemblies suffered a quality decline of 0.47% nucleotide identity when switching to the more distant reference, Nanopore assemblies declined by 1.22%. This emphasizes the great importance of choosing a suitable reference in mapping-based assemblies of Nanopore data. Using the *L. virgatum* reference produced a consensus of almost comparable quality to the *de novo* Nanopore assembly, and superior to the Illumina assembly based on the more distant reference (see Table 6).

A second essential point for obtaining high-quality reference-based assemblies with Nanopore as well as Illumina data is the choice of software used for mapping and variant calling. Preliminary tests with popular short-read mappers (Bowtie2 [88], BWA-MEM [89], and BBMap) showed that despite all three tools claim to support gapped alignment, only BBMap (which also supports long reads) was able to correctly map reads spanning large gaps (for mapping against the *Artemisia* reference, the largest deletion occurring in *L. vulgare* had a length of 459 bp). Bowtie2 and BWA-MEM either soft-clipped the respective reads or did not map them at all, leaving the site of deletion with a coverage gap. However, correct mapping across deletions is of utter importance as most variant callers will call the reference sequence at positions where read information is lacking, which would distort the resulting consensus sequence. Tested consensus-calling pipelines also included BCFtools *mpileup* plus *call* for Illumina data, which however was not able to correctly call indels as variants. For Nanopore data, variant calling was only successful using *callvariants2.sh* after *mapPacBio.sh* mapping.

An advantage of reference-based assembly is that the assembly process can be easily monitored by comparing the visualized mapping (e.g., in IGV) to the final called consensus. Specific patterns produced by the used datatype or algorithm can thus be taken into account and potential errors avoided. For example, when mapping Nanopore reads, especially insertions will often not be mapped at a precise location, but slightly shifted up- or downstream in several reads, thus blurring the signal for *callvariants2.sh*, which will ultimately call the reference in these cases. This problem is aggravated with more distant references as exact alignment gets more difficult. Regarding the variant-calling process, mismatches and small (1-bp or 2-bp) indels were usually captured correctly in both Illumina and Nanopore mappings. By contrast, larger insertions in *L. vulgare* with respect to the reference were more likely to be ignored, leading to the large number of deletions observed in the consensus sequences (56.2-89.4% of all mismatches, see Table 6). This effect seems to be owed to the used algorithm, intensifies with insertion length as well as more distant references, and is generally more pronounced with Nanopore data due to blurred mapping as described above. Deletions in the mapped taxon with respect to the reference were however more often called correctly from Nanopore reads, and almost always from Illumina reads. *Callvariants2.sh* also deals well with declining read coverage in homopolymer runs and mostly makes the correct full-length call.

### 4.6. Optimizing Nanopore *de novo* and reference-based assembly

A read coverage of at least 30-fold is generally recommended for Illumina *de novo* as well as reference-based plastome assembly [64, 6]. In our Illumina *de novo* assembly of *L. vulgare*, only 0.26% of bases received coverage below that threshold (Table 3). Accordingly, the Unicycler pipeline yielded the lowest possible number of contigs when using short reads and no read extension methodology [6]. The *L. virgatum* assembly, with 6.62% of bases having fewer than 30-fold coverage, resulted in five contigs. Although a complete plastome sequence was easily obtained, this somehow suboptimal result might be due to a lowered k-mer frequency as a consequence of low coverage in certain regions. Regarding Nanopore *de novo* assembly, a coverage of 30-fold will likely not yield optimal results (but see [11]). However, a minimum of 50-fold coverage in over 99.5% of the bases seemed to be sufficient for reliable assembly (Table 3). If coverage is too low, problems might arise with reads derived from PCR fragments with only small initial overlap; after adapter-, primer-, and quality-trimming, too many reads might get too short, precluding a continuous assembly. Fragment 9 and 10 in our analyses had an overlap of only 134 bp between the two primer sequences; this resulted in a very low coverage of down to 5-fold in the Nanopore-only assembly, and even a gap in a preliminary run with different settings (not shown). In Nanopore mapping assemblies, very low coverage might result in variants not being called, which will compromise the resulting consensus.

However, a much more important problem observed with Nanopore data is the detrimental effect of too high coverage. Wang et al. (2018) recognized an effect of too low or high coverage on assembly quality, stating that “there is a complex relationship between assembly accuracy and input read coverage when assembling chloroplast genomes with ONT data using Canu” [21]. [6] mention that *de novo* assembly performance could be improved by downsampling the input reads, and [86] stress that very high coverage may result in alternative assemblies which can lead to contig fragmentation. For the present study, preliminary *de novo* assembly tests (results not shown) with a reduced compared to the full dataset considerably decreased the number of contigs (approximately by half). The observed effect is likely due to the fact that at very high coverage, erroneous variants introduced by PCR or sequencing errors will be frequent enough in a dataset to pass the threshold for being incorporated into (extra) contigs by the assembler. To avoid this, a coverage-reduced dataset should be used for *de novo* assembly, for example via the *readSamplingBias* parameter in Canu. This has the side-effect of preferentially using the longest reads for analysis, which might also help to improve the results. With long-range PCR based reads, however, care must be taken to ensure that reads originating from short fragments are not accidentally filtered completely.

Filtering plastome reads from possible nuclear or contaminant reads, for example by mapping to a known reference, is a common procedure prior to assembly. However, applying too stringent filters might result in the loss of true plastid reads in the case of atypical plastomes or cp genomes that have incorporated mitochondrial or nuclear DNA [6]. For this reason, read filtering other than removing the PhiX spike-in was omitted in Illumina *de novo* analyses shown here. Unicycler proved relatively robust to this kind of perturbation: apparently, present contamination reads in the *L. vulgare* sample were assembled into an additional contig (showing BLAST matches in Proteobacteria) that could simply be excluded from further processing. In contrast to Unicycler, Canu was much more susceptible to contaminant-related errors. Even though input reads were subjected to BLAST filtering prior to *de novo* assembly, the latter still yielded a chimeric contig with one large part matching the *Artemisia* chloroplast and a small part blasting within Proteobacteria. Although the removal of this contig did not deteriorate the quality of the final assembly, the risk of obtaining erroneous contigs might be higher with Canu.

Another severe problem with the Nanopore reads used here for *de novo* assembly was the occurrence of chimeric reads. Two types of chimeras were observed. The first comprised reads with a first part being identical to a second part, but in reverse-complement orientation (so-called palindromes). It remains unclear what the source of this kind of error was. [47] describe a similar phenomenon in long reads that can occur during whole-genome amplification, so palindrome formation in cycles of long-range PCR at least seems conceivable. In contrast to the short, fragmented Illumina reads, the long Nanopore reads will introduce these errors into assembly. To avoid this, before starting *de novo* assembly we subjected reads to Pacasus, which splits reads containing palindromic sequences. Splitting proved essential, as initial assemblies with uncorrected reads produced a large number of (sometimes palindromic) contigs, rendering impossible the generation of a proper consensus sequence.

According to [90], who analyzed several types of chimeric reads originating from MinION sequencing, these can also consist of dissimilar fragments which are concatenated by the same sequencing adapter accidentally ligated in between them during library preparation. This type of chimera might also result from a very rare, Nanopore-specific process which has been called “*in silico* chimerism” by [90]. Here, too fast pore reloading results in the recognition of two fragments as one by the base-calling algorithm. A corresponding filtering step is incorporated in the Nanopore bioinformatic pipeline (Porechop). In the analyses presented here, 22.6% of the total flow cell reads were discarded due to putative middle adapters. Although the filtering was set to be quite stringent, it seems possible that chimeric reads passed the filter, and acted as bridges to connect sequence regions in Canu contigs which are actually distant from each other on the plastome. Indeed, from the ten total contigs produced by Canu for Nanopore and hybrid assemblies, three had to be split due to such effects. It must be stressed here that without a reference plastome, detection of chimeric contigs would not have been possible, and this technique of course harbors the risk of missing true rearrangements in the plastome to be assembled.

As a concluding remark, the frequency of chimeras in our dataset seems to be unusually high, considering the fact that all types of chimeras found by [90] together comprised only 1.7% of the reads and [47] described palindromic reads as occurring during whole-genome amplification, not long-range PCR. Other, unknown processes may thus be responsible for the phenomena observed. At least, both contaminant as well as chimeric reads are negligible problems for reference-based assemblies, as the initial mapping step will result in dismissal of the former as unmapped, while the latter will likely result in supplementary mappings, which are also excluded from the output if BBMap is used with the settings described here.

### 4.7. Conclusions - Lessons learned

Based on the results obtained in this study, we provide some rough guidelines on the sequencing of plant cp genomes. Regarding data production, PCR-mediated genome reduction has the advantage of being easy to realize in an ordinary lab, but will depend on the performance of universal primer sets in the studied group. Several shortcomings must also be taken into account (see section 4.1.). For enrichment of plastid DNA with subsequent Nanopore sequencing, the use of PCR must be regarded as suboptimal as it nullifies some of the main advantages of the latter. While Nanopore sequencing can now produce reads with lengths of more than two Mb [91], PCR-derived reads will currently have a maximum length of 23 kb [60]. This also precludes the disambiguation of the two IRs, which may otherwise be possible. If the use of long-range PCR plus Nanopore sequencing is intended, it is advisable to

- use a high-quality, proofreading polymerase for minimal entry of PCR artifacts into reads;
- choose fragment sizes as large as possible; this also requires the use of appropriate DNA extraction techniques;
- avoid too short overlaps among PCR fragments (preferably, > 300 bp) as in combination with low Nanopore read coverage, these might lead to fragmented assemblies;
- downsample regions of uneven coverage, for example fragment overlaps with doubled coverage, using VariantBam [92] or related tools; and
- account for palindromic sequences which might occur whenever PCR is used; these must be properly handled prior to assembly with the Pacasus tool or other appropriate software.

Hybrid assembly approaches with both Nanopore and Illumina data still yield better results than only using Nanopore reads. However, depending on the research question, Nanopore sequencing is also capable of delivering high-quality results at a comparatively lower cost. Our results show that the vast majority of the 221 deletion errors within the Nanopore *de novo* assembly lies within homopolymer regions of five or more bases; the same is true for a number of insertion errors. Exclusion of such regions will greatly reduce the error rate of the respective assembly. If phylogenetic analysis of several cp genomes in a plant group is intended, the genomes can be easily sequenced and analyzed after masking the respective homopolymer regions, which should be excluded from this type of analyses anyway. Of course, the degree of relatedness in the respective plant group must be considered. While in the Nanopore *de novo* assembly, the error rate of 0.41% sequence identity (including all errors) is much lower than the average divergence between *A. frigida* and *L. vulgare* (3.69%), the situation looks somewhat different in a close-knit species group like the genus *Leucanthemum* (*L. vulgare* - *L. virgatum*: 0.83%).

To ensure that high-quality assemblies can be obtained from analyzing Nanopore data alone, several important points should be taken into account:

- reads with middle adapters (i.e., chimeras consisting of two different reads) must be rigorously removed or at least bioinformatically split, to avoid the introduction of wrong proximity information into assembly
- for *de novo* assembly using Canu, the elimination of contaminant reads (nuclear DNA, other organisms) prior to assembly is vital. This can be done by blasting against a curated database of known plant genomes
- read coverage should be kept at reasonable heights for *de novo* assembly, especially when using uncorrected reads (as a gross starting point, not higher than 300-fold for the final unitigging step), for example by subsampling reads before assembly (e.g., with SAMtools or BBTools) or by applying the Canu parameters *readSamplingBias* / *readSamplingCoverage*
- preferentially including the longest reads might improve assembly quality; use the above Canu parameters or the *minReadLength* parameter in Canu

In some situations, for example with limited computational resources or large amounts of samples to assemble, reference-based assembly might be favored over *de novo* approaches. However, it should not be conducted in groups with known plastome variability or structural reorganization, but rather for population genomics or in genera of very closely related species. If these requirements are met, mapping assemblies can yield a consensus comparable to that of *de novo* assembly, as shown in the present study. For this, it is essential to

- use suitable software, for example BBMap plus the *callvariants2.sh* script. However, discrimination of the two IR copies will not be possible with BBMap as it cannot handle very long reads;
- correctly set software parameters. This may require some optimization; for a new plant group, it might be advisable to use parallel Illumina sequencing in the first plastome. for example, variant calling based on a majority vote for alleles might not work for the error-prone Nanopore reads. The minimum allele fraction required for a call could therefore be lowered (e.g., to 0.45); and
- choose the closest reference available (from the same genus, ideally the same species). According to our results, the quality of Nanopore reference-based assemblies decreases disproportionally fast with more divergent reference genomes, even if these still have sequence identities above 95% to the studied taxon. The reference should include the complete IR_A_ or only parts of it, depending on read length, to allow unambiguous mapping of all reads.

## Acknowledgements

The authors wish to thank Gunter Meister and Norbert Eichner at the Department of Biochemistry I (RNA-Biology) at the University of Regensburg, for kindly providing access to their Illumina sequencing facility and Tapestation device, and for helpful advice regarding library preparation and sequencing. We are also grateful to Dina Grohmann and Felix Grünberger from the Department of Microbiology (Archaea center) for providing equipment and laboratory material needed for Nanopore sequencing, and for kind help regarding Nanopore data processing and analysis. Michael Rehli and Alexander Fischer (Laboratory for Mononuclear Phagocyte Biology at the University Medical Center Regensburg) are acknowledged for spike-in of two library samples into one of their MiSeq sequencing runs and helpful suggestions regarding size selection. Special thanks are due to Ulrich Lautenschlager and Tankred Ott for their patient and constant help regarding all aspects of bioinformatic analyses, and for providing scripts and useful advice. We thank Anja Heuschneider for technical assistance and invaluable help in the lab. Susann Wicke is acknowledged for her kind advice regarding chloroplast genome annotation.

## Supporting Information Captions

**S1 Table. Taxa analyzed in the present study, alongside locality and collector information.**

**S2 Table. Primers used in the present study.** Primers for long-range PCR are based on those in Uribe-Convers et al. (2014) [28] and result in 16 chloroplast fragments. For fragments marked with an *, one or both primers were replaced by other primers published in Uribe-Convers et al. (2014). Modified primer sequences are indicated by the name suffix “_mod” or “-rc_mod”. For PCR and subsequent Sanger sequencing of the single-copy/inverted repeat junctions, five new primers were designed (given in bold type), the others were taken from Uribe-Convers et al. (2014). The gap between fragments 14 and 15 was likewise closed by Sanger sequencing of a PCR fragment based on newly designed primers.

**S3 Fig. Chloroplast genome map for *Leucanthemum virgatum*.** Genes on the outside of the outer circle are transcribed counterclockwise, genes on the inside are transcribed clockwise. Introns are illustrated with white color within genes; genes containing an intron are additionally marked with *. Pseudogenes are preceded by a ψ. The trans-spliced *rps12* gene is marked with °. Color-coding of genes depicts their affiliation to the functional groups given. The inner circle indicates the borders of the large single-copy (LSC) and small single-copy (SSC) regions as well as the inverted repeats (IR). The innermost gray shaded area shows the G+C content of the cp genome. The gene order is identical in *L. vulgare* (see Fig 2), whereas their exact positions and the extent of the inverted repeat slightly differ (Fig 3).

**S4 Table. Plastome characteristics for three Anthemideae species, and sequence divergence compared to *Leucanthemum vulgare*.** Length, GC content and gene information for *L. vulgare* and *L. virgatum* as obtained in the present study based on the Illumina *de novo* assemblies; information for *A. frigida* as taken from Liu et al. (2013) [71]. Percentages of identity and ‘alignment positions with N’ are based on the alignment length of the respective species and *L. vulgare*; ‘alignment positions with N’ denotes the amount of Ns required for alignment to *L. vulgare*. *L.*, *Leucanthemum*; bp, basepairs; IR, inverted repeat; n.a., not applicable; mism., mismatches.

**S5 File. Consensus sequences of comparative assemblies generated in the present study for *Leucanthemum vulgare*, excluding Illumina *de novo* assemblies.** Mapping assemblies given without the second inverted repeat (IRa). Nanopore *de novo* assembly (Lvulgare_Nanopore_denovo.fasta); hybrid *de novo* assembly of Illumina + Nanopore data (Lvulgare_Nanopore_hybrid_denovo.fasta); Illumina mapping assembly using *L. virgatum* reference (Lvulgare_Illumina_mapping_virgatum_oIR2.fasta); Illumina mapping assembly using *Artemisia frigida* reference (Lvulgare_Illumina_mapping_Artemisia_oIR2.fasta); Nanopore mapping assembly using *L. virgatum* reference (Lvulgare_Nanopore_mapping_virgatum_oIR2.fasta); Nanopore mapping assembly using *A. frigida* reference (Lvulgare_Nanopore_mapping_Artemisia_oIR2.fasta).

## References

1. Wicke S, Schneeweiss GM, Claude W, dePamphilis KF, Quandt MD. The evolution of the plastid chromosome in land plants: gene content, gene order, gene function. Plant Mol Biol. 2011; 76: 273–297.

2. Wicke S, Schneeweiss GM. Next-generation organellar genomics: potentials and pitfalls of high-throughput technologies for molecular evolutionary studies and plant systematics. In: Hörandl E, Appelhans MS, editors. Next Generation Sequencing in Plant Systematics, Regnum vegetabile vol. 158. Oberreifenberg: Koeltz Botanical Books; 2015. pp. 9–50.

3. Bendich AJ. Circular chloroplast chromosomes: the grand illusion. Plant Cell. 2014; 16: 1661–1666.

4. Kolodner R, Tewari KK. Inverted repeats in chloroplast DNA from higher plants. Proc Natl Acad Sci U S A. 1979; 76: 41–45.

5. Wolfe KH, Li W-H, Sharp PM. Rates of nucleotide substitution vary greatly among plant mitochondrial, chloroplast, and nuclear DNAs. Proc Natl Acad Sci U S A. 1987; 84: 9054–9058.

6. Twyford AD, Ness RW. Strategies for complete plastid genome sequencing. Mol Ecol Resour. 2017; 17: 858–868.

7. Laehnemann D, Borkhardt A, McHardy AC. Denoising DNA deep sequencing data-high-throughput sequencing errors and their correction. Brief Bioinform. 2016; 17: 154–179.

8. Bleidorn C. Third generation sequencing: technology and its potential impact on evolutionary biodiversity research. Syst Biodivers. 2016; 14: 1–8.

9. Rhoads A, Au KF. PacBio sequencing and its applications. Genomics Proteomics Bioinformatics 2015; 13: 278–289.

10. Rang FJ, Kloosterman WP, de Ridder J. From squiggle to basepair: computational approaches for improving nanopore sequencing read accuracy. Genome Biol. 2018; 19: 90.

11. Loman NJ, Quick J, Simpson JT. A complete bacterial genome assembled de novo using only nanopore sequencing data. Nat Methods. 2015; 12: 733–735.

12. Jain M, Olsen HE, Paten B, Akeson M. The Oxford Nanopore MinION: delivery of nanopore sequencing to the genomics community. Genome Biol. 2016; 17: 239.

13. Koren S, Schatz MC, Walenz, BP, Martin J, Howard JT, Ganapathy G, et al. Hybrid error correction and de novo assembly of single-molecule sequencing reads. Nat Biotechnol. 2012; 30: 693–700.

14. Belser C, Istace B, Denis E, Dubarry M, Baurens FC, Falentin C, et al. Chromosome-scale assemblies of plant genomes using nanopore long reads and optical maps. Nat Plants. 2018; 4: 879–887.

15. Gao K, Li J, Khan WU, Zhao TY, Yang X, Yang XY, et al. Comparative genomic and phylogenetic analyses of Populus section Leuce using complete chloroplast genome sequences. Tree Genet Genomes. 2019; 15: 32.

16. Kang H-I, Lee HO, Lee IH, Kim IS, Lee S-W, Yang TJ, et al. Complete chloroplast genome of Pinus densiflora Siebold & Zucc. and comparative analysis with five pine trees. Forests. 2019; 10: 600.

17. Chin CS, Alexander DH, Marks P, Klammer AA, Drake J, Heiner C, et al. Nonhybrid, finished microbial genomes assemblies from long-read SMRT sequencing data. Nat Methods. 2013; 10: 563–569.

18. Chaney L, Mangelson R, Ramaraj T, Jellen EN, Maughan PJ. The complete chloroplast genome sequences for four *Amaranthus* species (Amaranthaceae). Appl Plant Sci. 2016; 4: 1600063.

19. Xiang BB, Li XX, Qian J, Wang LZ, Ma L, Tian XX, et al. The complete chloroplast genome sequence of the medicinal plant *Swertia mussotii* using the PacBio RS II platform. Molecules. 2016; 21: 1029.

20. Ferrarini M, Moretto M, Ward JA, Šurbanovski N, Stevanović V, Giongo L, et al. An evaluation of the PacBio RS platform for sequencing and de novo assembly of a chloroplast genome. BMC Genomics. 2013; 14: 670.

21. Wang WW, Schalamun M, Morales-Suarez A, Kainer D, Schwessinger B, Lanfear R. Assembly of chloroplast genomes with long- and short-read data: a comparison of approaches using *Eucalyptus pauciflora* as a test case. BMC Genomics. 2018; 19: 977.

22. Leggett RM, Clark MD. A world of opportunities with nanopore sequencing. J Exp Bot. 2017; 68: 5419–5429.

23. Jansen HJ, Liem M, Jong-Raadsen SA, Dufour S, Weltzien F-A, Swinkels W, et al. Rapid de novo assembly of the European eel genome from nanopore sequencing reads. Sci Rep. 2017; 7: 7213.

24. Bethune K, Mariac C, Couderc M, Scarcelli N, Santoni S, Ardisson M, et al. Long-fragment targeted capture for long-read sequencing of plastomes. Appl Plant Sci. 2019; 7: e1243.

25. Oberprieler C, Himmelreich S, Vogt R. A new subtribal classification of the tribe Anthemideae (Compositae). Willdenowia. 2007; 37: 89–114.

26. Doyle JJ, Doyle JS. A rapid DNA isolation procedure for small quantities of fresh leaf tissue. Phytochem Bull. 1987; 19: 11–15.

27. Doyle JJ, Dickson EE. Preservation of plant samples for DNA restriction endonuclease analysis. Taxon. 1987; 36: 715–722.

28. Uribe-Convers S, Duke JR, Moore MJ, Tank DC. A long PCR–based approach for DNA enrichment prior to next-generation sequencing for systematic studies. Appl Plant Sci. 2014; 2: 1300063.

29. Morgulis A, Coulouris G, Raytselis Y, Madden TL, Agarwala R, Schäffer AA. Database indexing for production MegaBLAST searches. Bioinformatics. 2008; 15: 1757–1764.

30. Johnson M, Zaretskaya I, Raytselis Y, Merezhuk Y, McGinnis S, Madden TL. NCBI BLAST: a better web interface. Nucleic Acids Res. 2008; 36: W5–W9.

31. Jansen RK, Palmer JD. A chloroplast DNA inversion marks an ancient evolutionary split in the sunflower family (Asteraceae). Proc Natl Acad Sci U S A. 1987; 84: 5818–5822.

32. Kim K-J, Choi K-S, Jansen RK. Two chloroplast inversions originated simultaneously during the early evolution of the Sunflower family (Asteraceae). Mol Biol Evol. 2005; 22: 1783–1792.

33. Kück U, Bunse A, Holländer-Czytko H, Jeske S, Klämbt C, Klapper R, et al. Praktikum der Molekulargenetik. Berlin, Heidelberg. Springer; 2005. pp. 376–377.

34. Gremme G, Steinbiss S, Kurtz S. GenomeTools: a comprehensive software library for efficient processing of structured genome annotations. IEEE/ACM Trans Comput Biol Bioinform. 2013; 10: 645–656.

35. Bushnell B. BBTools software package. 2014. [cited 21 March 2019] http://sourceforge.net/projects/bbmap.

36. Andrews S, Lindenbaum P, Howard B, Ewels P. FastQC: a quality control tool for high throughput sequence data. 2010. [cited 21 March 2019] Available from: http://www.bioinformatics.babraham.ac.uk/projects/fastqc.

37. Li H, Handsaker B, Wysoker A, Fennell T, Ruan J, Homer N, et al. The Sequence alignment/map (SAM) format and SAMtools. Bioinformatics. 2009; 25: 2078–2079.

38. Okonechnikov K, Conesa A, García-Alcalde F. Qualimap 2: advanced multi-sample quality control for high-throughput sequencing data. Bioinformatics. 2015; 32: 292–294.

39. Robinson JT, Thorvaldsdóttir H, Winckler W, Guttman M, Lander ES, Getz G, et al. Integrative Genomics Viewer. Nat Biotechnol. 2011; 29: 24–26.

40. Danecek P, Auton A, Abecasis G, Albers CA, Banks E, DePristo MA, et al. The variant call format and VCFtools. Bioinformatics. 2011; 27: 2156–2158.

41. Wick RR, Judd LM, Gorrie CL, Holt KE. Unicycler: resolving bacterial genome assemblies from short and long sequencing reads. PLoS Comput Biol. 2017; 13: e1005595.

42. Bankevich A, Nurk S, Antipov D, Gurevich AA, Dvorkin M, Kulikov AS, et al. SPAdes: a new genome assembly algorithm and its applications to single-cell sequencing. J Comput Biol. 2012; 19: 455–477.

43. Walker BJ, Abeel T, Shea T, Priest M, Abouelliel A, Sakthikumar S, et al. Pilon: an integrated tool for comprehensive microbial variant detection and genome assembly improvement. PLoS One. 2014; 9: e112963.

44. Wick RR, Schultz MB, Zobel J, Holt KE. Bandage: interactive visualization of de novo genome assemblies. Bioinformatics. 2015; 31: 3350–3352.

45. de Coster W, d’Hert S, Schultz DT, Cruts M, van Broeckhoven C. NanoPack: visualizing and processing long-read sequencing data. Bioinformatics. 2018; 34: 2666–2669.

46. Camacho C, Coulouris G, Avagyan V, Ma N, Papadopoulos J, Bealer K, et al. BLAST+: architecture and applications. BMC Bioinformatics. 2008; 10: 421.

47. Warris S, Schijlen E, van de Geest H, Vegesna R, Hesselink T, Hekkert BTL, et al. Correcting palindromes in long reads after whole-genome amplification. BMC Genomics. 2018; 19: 798.

48. Koren S, Walenz BP, Berlin K, Miller JR, Bergmann NH, Phillippy AM. Canu: scalable and accurate long-read assembly via adaptive k-mer weighting and repeat separation. Genome Res. 2017; 27: 722–736.

49. Slater GS, Birney E. Automated generation of heuristics for biological sequence comparison. BMC Bioinformatics. 2005; 6: 31.

50. Li H. Minimap2: pairwise alignment for nucleotide sequences. Bioinformatics. 2018; 34: 3094–3100.

51. Hall TA. BioEdit: a user-friendly biological sequence alignment editor and analysis program for Windows 95/98/NT. Nucleic Acids Symp Ser. 1999; 41: 95–98.

52. Goodwin S, Gurtowski J, Ethe-Sayers S, Deshpande P, Schatz MC, McCombie WR. Oxford Nanopore sequencing, hybrid error correction, and de novo assembly of a eukaryotic genome. Genome Res. 2015; 25: 1750–1756.

53. Tillich M, Lehwark P, Pellizzer T, Ulbricht-Jones ES, Fischer A, Bock R et al. GeSeq – versatile and accurate annotation of organelle genomes. Nucleic Acids Res. 2017; 45: W6–W11.

54. Kent WJ. 2002. BLAT - The BLAST-like alignment tool. Genome Res. 12: 656–664.

55. Laslett D, Canback B. ARAGORN, a program to detect tRNA genes and tmRNA genes in nucleotide sequences. Nucleic Acids Res. 2004; 32: 11–16.

56. Lohse M, Drechsel O, Kahlau S, Bock R. OrganellarGenomeDRAW - a suite of tools for generating physical maps of plastid and mitochondrial genomes and visualizing expression data sets. Nucleic Acids Res. 2013; 41: W575–W581.

57. Greiner S, Lehwark P and Bock R (2019) OrganellarGenomeDRAW (OGDRAW) version 1.3.1: expanded toolkit for the graphical visualization of organellar genomes. Nucleic Acids Research, Web Server issue, DOI: dx.doi.org/10.1093/nar/gkz238.

58. Katoh K, Misawa K, Kuma K, Miyata T. MAFFT: a novel method for rapid multiple sequence alignment based on fast Fourier transform. Nucleic Acids Res. 2002; 30: 3059–3066.

59. Katoh K, Standley DM. MAFFT Multiple sequence alignment software version 7: improvements in performance and usability. Mol Biol Evol. 2013; 30: 772–780.

60. Yang JB, Li DZ, Li HT. Highly effective sequencing whole chloroplast genomes of angiosperms by nine novel universal primer pairs. Mol Ecol Resour. 2014; 14: 1024–1031.

61. Bakker FT. Herbarium genomics: skimming and plastomics from archival specimens. Webbia. 2017; 72: 35–45.

62. Rabah SO, Shrestha B, Hajrah NH, Sabir MJ, Alharby HF, Sabir Mernan, et al. *Passiflora* plastome sequencing reveals widespread genomic rearrangements. J Syst Evol. 2019; 57: 1–14.

63. Doorduin L, Gravendeel B, Lammers Y, Ariyurek Y, Chin-A-Woeng T, Vrieling K. The complete chloroplast genome of 17 individuals of pest species *Jacobaea vulgaris*: SNPs, microsatellites and barcoding markers for population and phylogenetic studies. DNA Res. 2011; 18: 93–105.

64. Straub SC, Parks M, Weitemier K, Fishbein M, Cronn RC, Liston A. Navigating the tip of the genomic iceberg: Next generation sequencing for plant systematics. Am J Bot. 2012; 99: 349–364.

65. Cronn R, Knaus BJ, Liston A, Maughan PJ, Parks M, Syring JV, et al. Targeted enrichment strategies for next-generation plant biology. Am J Bot. 2012; 99: 291–311.

66. Mariac C, Scarcelli N, Pouzadou J, Barnaud A, Billot C, Faye A, et al. Cost-effective enrichment hybridization capture of chloroplast genomes at deep multiplexing levels for population genetics and phylogeography studies. Mol Ecol Res. 2014; 14: 1103–1113.

67. Takamatsu T, Baslam M, Inomata T, Oikawa K, Itoh K, Ohnishi T, et al. Optimized method of extracting rice chloroplast DNA for high-quality plastome resequencing and de novo assembly. Front Plant Sci. 2018; 9: 266.

68. Civáň P, Foster PG, Embley MT, Séneca A, Cox CJ. Analyses of charophyte chloroplast genomes help characterize the ancestral chloroplast genome of land plants. Genome Biol Evol. 2014; 6: 897–911.

69. Konowalik K, Wagner F, Tomasello S, Vogt R, Oberprieler C. Detecting reticulate relationships among diploid *Leucanthemum* Mill. (Compositae, Anthemideae) taxa using multilocus species tree reconstruction methods and AFLP fingerprinting. Mol Biol Evol. 2015; 92: 308–328.

70. Wagner F, Ott T, Zimmer C, Reichhart V, Vogt R, Oberprieler C. ‘At the crossroads towards polyploidy’: genomic divergence and extent of homoploid hybridization are drivers for the formation of the ox-eye daisy polyploid complex (*Leucanthemum*, Compositae-Anthemideae). New Phytol. 2019; 223: 2039–2053.

71. Liu Y, Huo N, Dong L, Wang Y, Zhang S, Young HA, et al. Complete chloroplast genome sequences of Mongolia medicine *Artemisia frigida* and phylogenetic relationships with other plants. PLoS One. 2013; 8: e57533.

72. Walker JF, Jansen RK, Zanis MJ, Emery NC. Sources of inversion variation in the small single copy (SSC) region of chloroplast genomes. Am J Bot. 2015; 102: 1751–1752.

73. Palmer JD. Chloroplast DNA exists in two orientations. Nature. 1983; 301: 92–93.

74. Timme RE, Kuehl JV, Boore JL, Jansen RK. A comparative analysis of the *Lactuca* and *Helianthus* (Asteraceae) plastid genomes: identification of divergent regions and categorization of shared repeats. Am J Bot. 2007; 94: 302–312.

75. Shaw J, Shafer HL, Leonard OR, Kovach MJ, Schorr M, Morris AB. Chloroplast DNA sequence utility for the lowest phylogenetic and phylogeographic inferences in angiosperms: the tortoise and the hare IV. Am J Bot. 2014; 101: 1987–2004.

76. Curci PL, De Paola D, Danzi D, Vendramin GG, Sonnante G. Complete chloroplast genome of the multifunctional crop globe artichoke and comparison with other Asteraceae. PLoS One. 2015; 10: e0120589.

77. Dong W, Xu C, Li C, Sun J, Zuo Y, Shi S, et al. *ycf1*, the most promising plastid DNA barcode of land plants. Sci Rep. 2015; 5: 8348.

78. Zhu A-D, Guo W-H, Gupta S, Fan W-S, Mower JP. Evolutionary dynamics of the plastid inverted repeat: the effects of expansion, contraction, and loss on substitution rates. New Phytol. 2016; 209: 1747–1756.

79. Birky CW, Walsh JB. Biased gene conversion, copy number, and apparent mutation rate differences within chloroplast and bacterial genomes. Genetics. 1992; 130: 677–683.

80. Perry AS, Wolfe KH. Nucleotide substitution rates in legume chloroplast DNA depend on the presence of the inverted repeat. J Mol Evol. 2002; 55: 501–508.

81. Schmidt MHW, Vogel A, Denton AK, Istace B, Wormit A, van de Geest H, et al. De novo assembly of a new *Solanum pennellii* accession using Nanopore sequencing. Plant Cell. 2017; 29: 2336–2348.

82. Wang SB, Song QW, Li SS, Hu ZG, Dong GQ, Song C, et al. Assembly of a complete mitogenome of *Chrysanthemum nankingense* using Oxford Nanopore long reads and the diversity and evolution of Asteraceae mitogenomes. Genes. 2018; 9: 547.

83. Kolmogorov M, Yuan J, Lin Y, Pevzner P. Assembly of long, error-prone reads using repeat graphs. Nat Biotechnol. 2019; 37: 540–546.

84. Huang H, Shi C, Liu Y, Mao S-Y, Gao L-Z. Thirteen *Camellia* chloroplast genome sequences determined by high-throughput sequencing: genome structure and phylogenetic relationships. BMC Evol Biol. 2014; 14: 151.

85. Jansen RK, Raubeson LA, Boore JL, dePamphilis CD, Chumley TW, Haberle RC, et al. Methods for obtaining and analyzing whole chloroplast genome sequences. In: Zimmer EA, Roalson EH, editors. Methods in Enzymology vol. 395, Molecular Evolution: Producing the Biochemical data, part B. San Diego, London: Elsevier Academic Press; 2005. pp. 348–384.

86. Izan S, Esselink D, Visser RGF, Smulders MJM, Borm T. De novo assembly of complete chloroplast genomes from non-model species based on a k-mer frequency-based selection of chloroplast reads from total DNA sequences. Front Plant Sci. 2017; 8: 1271.

87. Sancho R, Cantalapiedra CP, López-Alvarez D, Gordon SP, Vogel JP, Catalán P et al. Comparative plastome genomics and phylogenomics of *Brachypodium*: flowering time signatures, introgression and recombination in recently diverged ecotypes. New Phytol. 2018; 218: 1631–1644.

88. Langmead B, Salzberg SL. Fast gapped-read alignment with Bowtie2. Nat Methods. 2012; 9: 357–359.

89. Li H, Durbin R. Fast and accurate short read alignment with Burrows-Wheeler Transform. Bioinformatics. 2009; 25: 1754–1760.

90. White R, Pellefigues C, Ronchese F, Lamiable O, Eccles D. Investigation of chimeric reads using the MinION [version 2; referees: 2 approved]. F1000Res. 2017; 6: 631.

91. Payne A, Holmes N, Rakyan V, Loose M. Whale watching with BulkVis: A graphical viewer for Oxford Nanopore bulk fast5 files. Bioinformatics. 2018; 35: 2193–2198.

92. Wala J, Zhang C-Z, Meyerson M, Beroukhim R. VariantBam: filtering and profiling of next-generational sequencing data using region-specific rules. Bioinformatics. 2016; 32: 2029–2031.

