## Supporting Information S1 Table for "Can we use it? On the utility of *de novo* and reference-based assembly of Nanopore data for plant plastome sequencing"

| Sample | Species | Label text and locality |
| --- | --- | --- |
| H3 | *Leucanthemum vulgare* Lam. | Germany, Regensburg, University, hill lawn in front of the faculty department of biology, near greenhouse, in culture at the botanical garden, 48°59’39’’N-12°05’30’’E, 09.01.2018, Anja Metko s.n. |
| A1221 | *Leucanthemum virgatum*  (Desr.) Clos | France, Tournefort, Val de la Tinée, Garrigue on calcareous rock, on the roadside, 225 m, 46°13'39.4968''N-2°12'49.4964''E, 30.05.2018, Davide Dagnino, Gabriele Casazza, Carmelo Macrì s.n. |
