## Supplementary figures and images for "Can we use it? On the utility of *de novo* and reference-based assembly of Nanopore data for plant plastome sequencing"

### Supporting Information S3 Fig

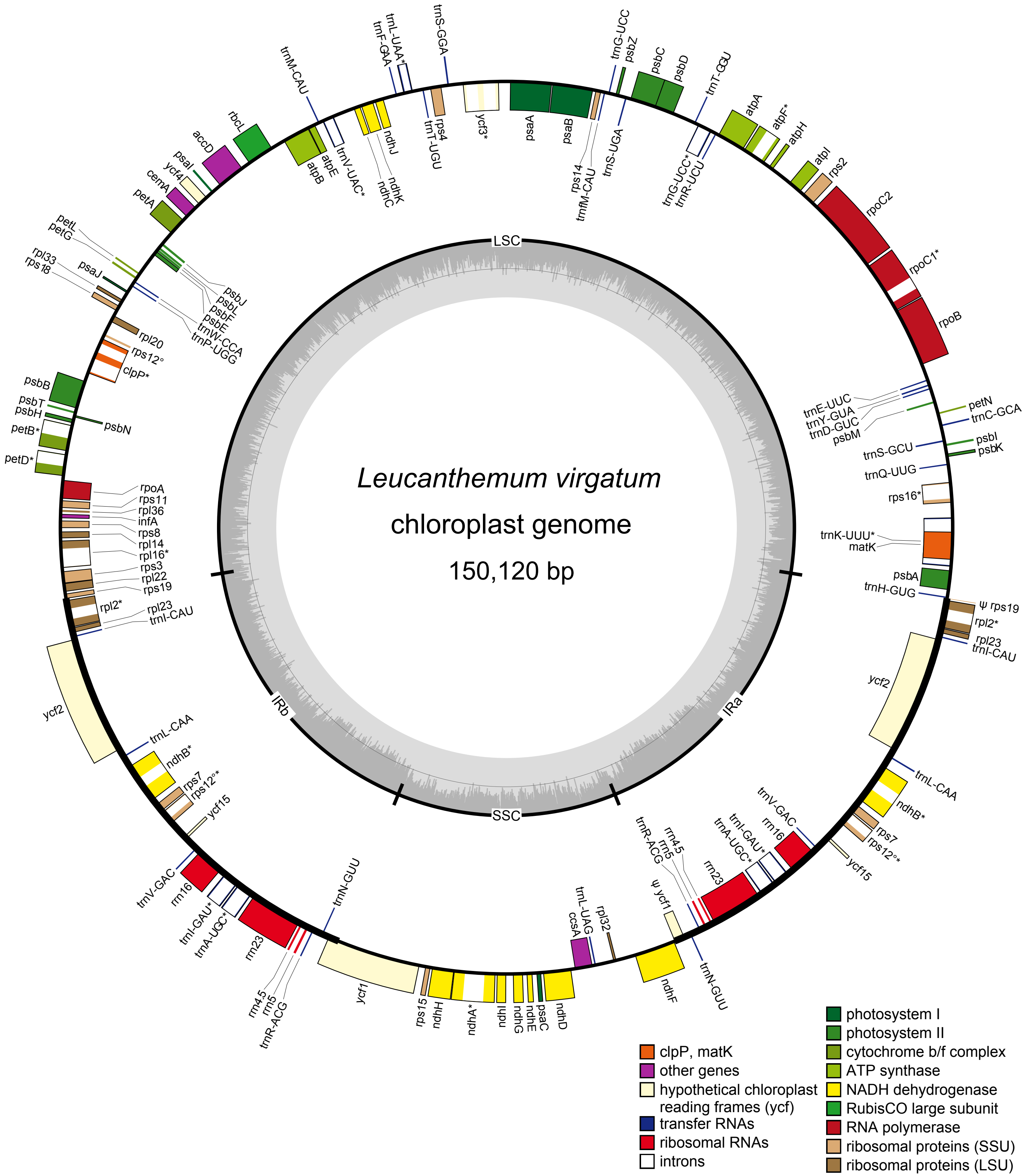
