## Supporting Information S4 Table for "Can we use it? On the utility of *de novo* and reference-based assembly of Nanopore data for plant plastome sequencing"

|  | *L. vulgare* | *L. virgatum* | *Artemisia frigida* |
| --- | --- | --- | --- |
| Genome size (bp) | 150,191 bp | 150,120 bp | 151,076 bp |
| LSC length (bp) | 82,675 bp | 82,641 bp | 82,740 bp |
| SSC length (bp) | 18,400 bp | 18,435 bp | 18,394 bp |
| IR length (bp) | 24,558 bp | 24,522 bp | 24,971 bp |
| % GC content | 37.45 | 37.45 | 37.48 |
| # of protein-coding genes | 80 | 80 | 80 |
| # of tRNA genes | 30 | 30 | 30 |
| # of rRNA genes | 4 | 4 | 4 |
| # of genes duplicated in IR | 18 | 18 | 18 |
| # of genes with introns | 18 | 18 | 18 |
| % identity to *L. vulgare* | n.a. | 99.17 | 96.31 |
| % alignment positions with N | n.a. | 0.003 | 0 |
| total mismatches | n.a. | 1,038 | 4,704 |
| no. of substitutions (% total mism.) | n.a. | 299 (28.8) | 1,725 (36.7) |
| no. of gaps (% total mismatches) | n.a. | 391 (37.7) | 1,272 (27.0) |
| deletion events | n.a. | 57 | 154 |
| no. of inserted bases (% total mism.) | n.a. | 348 (33.5) | 1,707 (36.3) |
| insertion events | n.a. | 64 | 136 |
